# Nutrient stress activates Rab5b-mediated autophagy to remodel the synaptic proteome

**DOI:** 10.1101/2025.11.25.690410

**Authors:** M Overhoff, L Ickert, S Marten, A Hill, F Tellkamp, SL Ludwig, P Antczak, FC Koehler, RU Müller, M Krüger, NL Kononenko

## Abstract

Synaptic proteostasis is crucial for maintaining neuronal function and plasticity, yet how synapses adapt to metabolic stress remains poorly understood. Here, we show that nutrient deprivation, particularly serum withdrawal, induces robust autophagy-dependent remodeling of the synaptic proteome, while mTORC1 inhibition has more limited effects. Nutrient stress rapidly activates autophagy both globally and at synapses, with synaptic autophagy peaking within 1–2 hours of serum withdrawal. Mechanistically, we uncover that the LC3 lipidation complex (ATG5–ATG12–ATG16L1) is recruited to synapses via Rab5b-positive endosomes in a dynein-dependent manner. Live imaging reveals enhanced Rab5b–ATG16L1 co-trafficking and increased ATG5 mobility upon serum withdrawal, supporting a model of spatiotemporally controlled autophagy precursor delivery to synaptic compartments. Functionally, nutrient deprivation acutely dampens neuronal excitability *in vitro*, while a two-week fasting-mimicking diet *in vivo* triggers synaptic proteome remodeling that overlaps with starvation-induced autophagy cargo. In contrast, restriction of mTORC1-activating amino acids fails to induce comparable synaptic changes, suggesting that synaptic autophagy is regulated by nutrient signals beyond mTORC1. Our findings define a Rab5b-mediated trafficking mechanism that couples nutrient sensing to localized synaptic degradation, providing new insight into how neurons preserve proteostasis under metabolic challenge.

## Introduction

Neurons rely on tightly regulated proteostasis mechanisms to maintain synaptic function and plasticity [1, 2]. At synapses, continuous remodeling of the proteome is essential for neuronal communication, learning, and memory [3–6]. Although synaptic proteins have an unusually long turnover [7–11], damaged proteins must be degraded, requiring precise coordination between protein synthesis, trafficking, and degradation to prevent dysfunction. Dysregulation of these processes has been implicated in neurodegenerative diseases, where impaired proteostasis leads to synaptic failure and neuronal loss [12, 13]. While autophagy has emerged as a key player in neuronal proteostasis [14–16], its role in synaptic protein homeostasis under conditions of metabolic stress remains poorly understood.

Nutrient availability is a fundamental regulator of cellular metabolism and survival [17], with neurons exhibiting unique adaptations to metabolic fluctuations [18–20]. One of the primary pathways governing metabolic adaptation is the mechanistic target of rapamycin complex 1 (mTORC1), a central regulator of cellular growth, autophagy, and protein synthesis [21]. Under nutrient-rich conditions, mTORC1 promotes anabolic processes while suppressing autophagy [22]. In contrast, nutrient deprivation inhibits mTORC1, triggering autophagy-mediated degradation of cellular components to maintain energy homeostasis. While the regulation of autophagy in neuronal cell bodies and growth cones has been extensively studied, how synapses integrate metabolic cues to modulate autophagy and proteome dynamics remains largely unexplored. Recent studies suggest that synaptic autophagy plays a critical role in maintaining synaptic integrity and plasticity [23], particularly in response to stress [24, 25]. However, the precise mechanisms through which metabolic stress influences autophagosome formation, trafficking, and cargo selection at synapses remain unclear. Moreover, whether mTORC1 inhibition and nutrient deprivation have distinct or overlapping effects on synaptic proteome remodeling has not been systematically examined.

In this study, we investigate the impact of nutrient deprivation and mTORC1 inhibition on synaptic proteome dynamics and autophagy induction. We demonstrate that the synaptic proteome exhibits differential vulnerability to metabolic stress, with distinct subsets of proteins affected by nutrient withdrawal and/or mTORC1 inhibition. We further show that serum deprivation is sufficient to trigger synaptic autophagy and enhance autophagosome trafficking, highlighting a dominant role for metabolic stress in synaptic proteostasis. Using *Atg5*-deficient neurons, we establish that nutrient deprivation-induced synaptic protein remodeling requires autophagy. Mechanistically, we reveal a temporally and spatially regulated recruitment of the LC3 lipidation system to synapses, as well as a selective interaction between autophagosomes and Rab5b that facilitates their synaptic localization using microtubule-based transport. Finally, we demonstrate that short-term serum starvation alters neuronal activity, whereas prolonged metabolic stress induces synaptic proteome remodeling *in vivo*.

Together, our findings provide mechanistic insights into how neurons adapt to metabolic stress via synaptic autophagy and proteome remodeling. Understanding these processes has important implications for neurodegenerative diseases, where metabolic dysfunction and impaired autophagy contribute to synaptic degeneration and neuronal loss.

## Results

### Differential vulnerability of the synaptic proteome to nutrient withdrawal and mTORC1 inhibition

To investigate how synaptic proteins respond to metabolic stress, we adapted an *ex vivo* protocol using acute cortical slices from adult mice. Slices were incubated for 6 hours in either control maintenance medium (see **Suppl. Table 1**), nutrient-deprived conditions—achieved by fetal calf serum (FCS) withdrawal or exposure to Earle’s Balanced Salt Solution (EBSS)—or with the mTORC1 inhibitor rapamycin (10 nM), as validated in **Fig. S1A,B**. Following treatment, synaptosomes were isolated (**Fig. S1C-E**) and subjected to mass spectrometry–based proteomic analysis (**Fig. 1A, Extended Table 1**). Comprehensive proteomic profiling revealed that each treatment elicited a unique pattern of synaptic protein regulation. Both serum withdrawal and EBSS treatment induced significant and widespread alterations in synaptic protein abundance (**Fig. 1B,F,H**), with a substantial number of proteins being markedly up- or downregulated compared to control. Serum withdrawal had the most pronounced effect, resulting in significant deregulation of 79 synaptic proteins (**Fig. 1D**). Gene ontology (GO) enrichment analysis confirmed that these proteins were predominantly associated with synaptic subcompartments (**Fig. 1E**). Notably, prominent synaptic components such as Cacnb1, Sh3gl1 (Endophilin A2), and Clta were upregulated upon serum deprivation, while proteins such as Grik5, Gabrb3, and Nrxn2 showed decreased levels under this condition (see **Fig. 1C**). While EBSS-induced changes were somewhat less extensive (**Fig. 1H**), this treatment triggered specific alterations in proteins involved in both synaptic function and cytoskeletal dynamics (**Fig. 1I**), suggesting convergent as well as divergent pathways of synaptic remodeling in response to different forms of nutrient stress. In contrast, rapamycin-mediated mTORC1 inhibition produced only minor changes in the overall synaptic proteome (**Fig. 1J–M**). GO-term enrichment analysis of rapamycin-sensitive proteins revealed enrichment in compartments such as the ER membrane and gamma-tubulin complex (**Fig. 1M**), highlighting remodeling of non-synaptic proteins under mTORC1 inhibition. Taken together, these data indicate that nutrient withdrawal, particularly serum removal, is a potent driver of synaptic proteome remodeling, while mTORC1 inhibition has a more selective effect, primarily targeting ER and cytoskeletal compartments. While some synaptic proteins were commonly regulated across conditions, the majority of changes were treatment-specific, underscoring mechanistically distinct responses to nutrient stress and mTORC1 inhibition. We note that serum withdrawal removes nutritional components and trophic factors simultaneously. Thus, compared to EBSS deprivation or selective mTORC1 inhibition, serum withdrawal represents a broader nutritional–trophic stress, likely contributing to its stronger effect on synaptic proteome remodeling.

**Figure 1.**
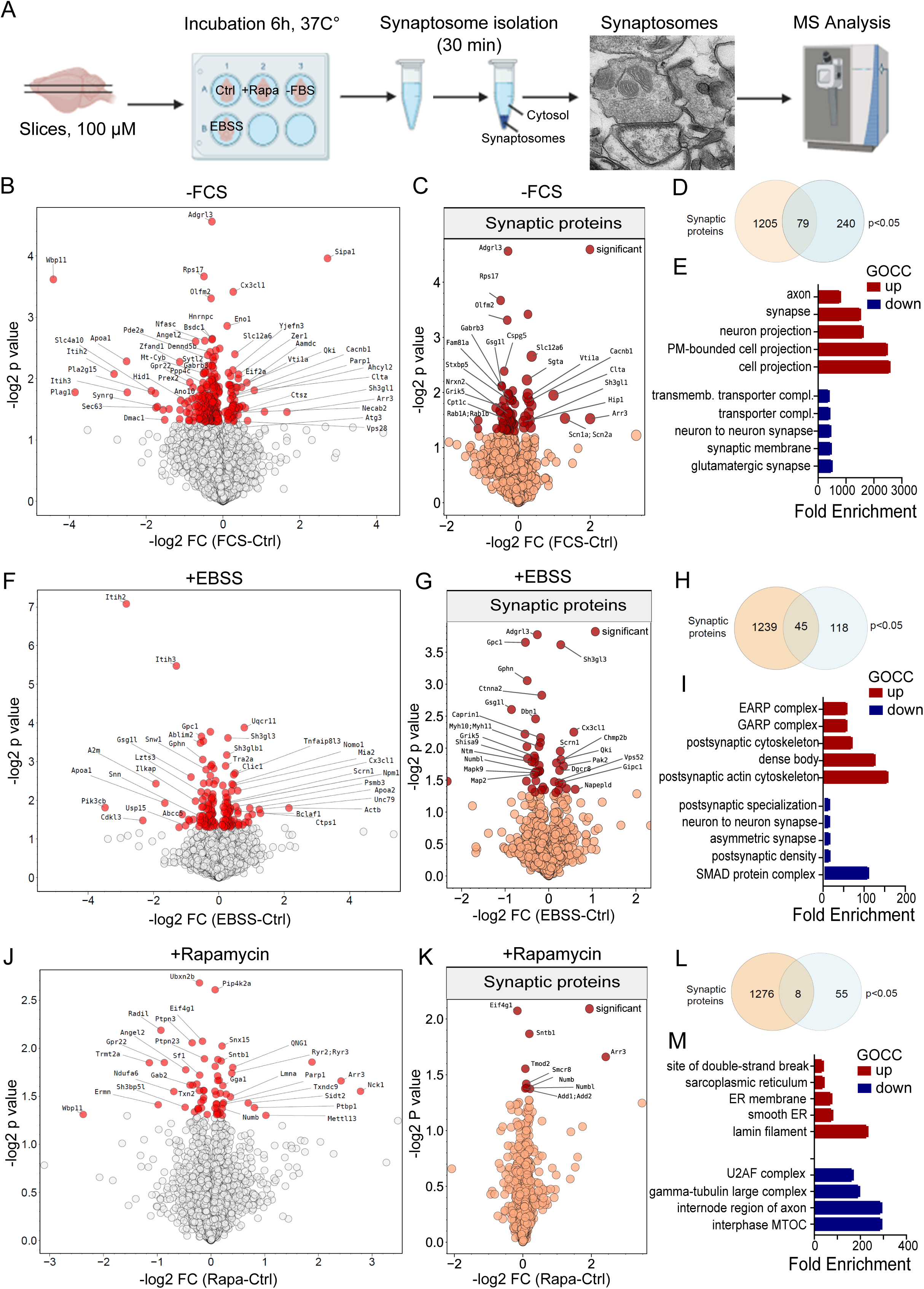
Differential remodeling of the synaptic proteome upon nutrient withdrawal and mTORC1 inhibition. **(A)** Schematic workflow of experimental design: acute cortical slices were either kept in control maintenance media or subjected to nutrient stress or mTORC1 inhibition (serum withdrawal, EBSS, or rapamycin (10 nM) treatment for 6h), followed by synaptosome isolation and label-free quantitative mass spectrometry (MS). (**B-E**) Differentially expressed proteins in synaptosomes upon 6h of serum starvation. Volcano plot showing differentially expressed total proteome (B), selectively deregulated synaptic proteins (C), their proportion in total deregulated proteome (D), and their GO term cellular component enrichment analysis (E). Selected proteins are annotated; significance is shown in red and reflects Student’s T-test at p<0.05. (**F-I**) Differentially expressed proteins in synaptosomes upon 6h of EBSS starvation. Volcano plot showing differentially expressed total proteome (G), selectively deregulated synaptic proteins (H), their proportion in total deregulated proteome (D), and their GO term cellular component enrichment analysis (I). Red colour in B and C highlights upregulated proteins at p < 0.05. (**J-M**) Differentially expressed proteins in synaptosomes upon 6h of rapamycin treatment. Volcano plot showing differentially expressed total proteome (J), selectively deregulated synaptic proteins (K), their proportion in total deregulated proteome (L), and their GO term cellular component enrichment analysis (M). Red colour in B and C highlights upregulated proteins at p < 0.05. Data represent n=5 biological replicates per condition.

### Autophagy is rapidly induced in cortical neurons and synapses upon nutrient deprivation

To determine whether metabolic stress induces neuronal autophagy and may contribute to the observed synaptic proteome remodeling, we first analyzed LC3 lipidation—a widely used marker for autophagosome formation—in primary cortical neurons cultured treated with serum-free medium (-FCS), EBSS, or rapamycin for 6 hours, in the presence or absence of the lysosomal inhibitor Bafilomycin A1 (BafA) (**Fig. 2A**). All treatments increased LC3-II levels in the presence of BafA, indicating enhanced autophagic flux (**Fig. 2B,C**). Notably, BafA alone elevated LC3-II levels even in control neurons, suggesting that basal autophagy is constitutively active under standard culture conditions, consistent with previous reports [26]. Rapamycin, serum withdrawal and EBSS, all modestly enhanced LC3-II accumulation with serum deprivation exerting the strongest effect. These findings were corroborated by stabilization of the autophagy cargo receptor p62 (**Fig. S2A,B**), further supporting modest but detectable increases in autophagic flux under nutrient stress.

**Figure 2.**
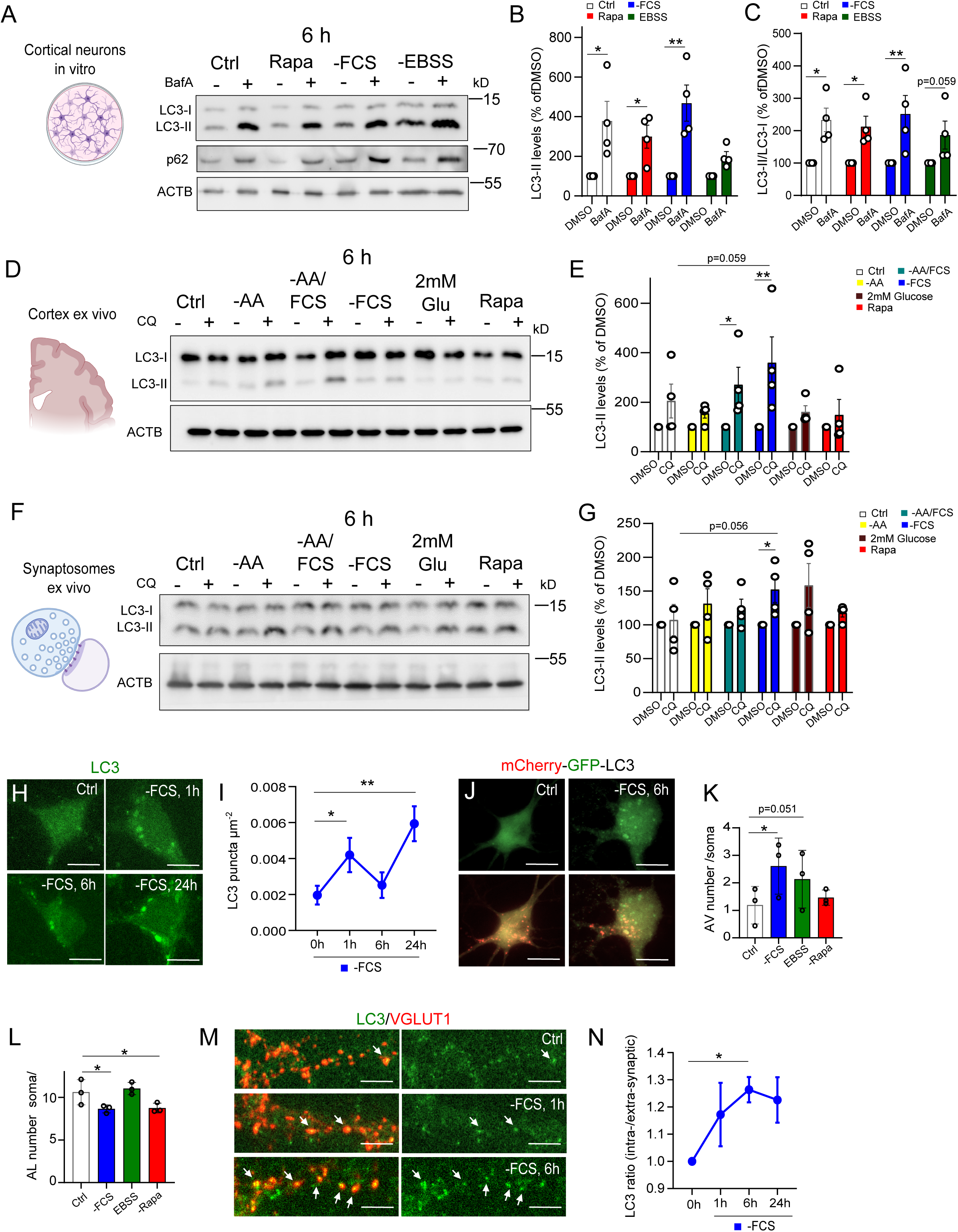
Rapid induction of autophagy in neurons and synapses upon nutrient stress. (**A-C**) Western blot analysis (A) and quantification of LC3-II levels (B) and/or LC3-II/LC3-I ratio (C) in primary cortical neurons treated for 6h with serum-free medium (-FCS), EBSS, rapamycin (10 nM) or kept in control maintenance medium in the presence or absence of 67 nM of Bafilomycin A1 (BafA) (n=3 of biological replicates). Baseline levels in DMSO-treated controls are normalized to 100%. (**D, E**) Western blot analysis (D) and quantification (E) of LC3-II levels in cytosolic fraction isolated from acute cortical slices treated for 6□h with amino acid-free medium (-AA), serum-free medium (-FCS), -AA/FCS-free medium, cultured medium containing 2mM glucose, rapamycin (10 nM) or kept in control medium in the presence or absence of 400 µM of chloroquine (CQ (n=4 of biological replicates). Baseline levels in DMSO-treated controls are normalized to 100%. (**F, G**) Western blot analysis (F) and quantification (G) of LC3-II levels in synaptosomal fraction isolated from acute cortical slices treated for 6 h with amino acid-free medium (-AA), serum-free medium (-FCS), -AA/FCS-free medium, cultured medium containing 2mM glucose, rapamycin (10 nM) or kept in control medium in the presence or absence of 400 µM of chloroquine (CQ) (n=4 of biological replicates). Baseline levels in DMSO-treated controls are normalized to 100%. (**H, I**) Immunofluorescence analysis of LC3 puncta in neuronal soma upon 1h, 6h and 24h of FCS withdrawal (100 neurons/timepoint from n=5 independent experiments). Scale bars: 5µm. (**J–L**) Live cell imaging (J) and quantification of autopjagic vesciels (AV; GFP^+^/mCherry^+^) (K) and autolysosomes (AL; GFP^−^/mCherry^+^) (L) in cultured cortical neurons expressing tandem-tagged mCherry-GFP-LC3 reporter after 2 h of nutrient deprivation (-FCS, EBSS or rapamycin (10 nM)) (25-32 neurons/condition from n=3 biological replicates). AV-autophagosome. Scale bars: 10µm. (**M,N**) Immunofluorescence analysis of LC3 levels at VGLUT1-positive synaptic sites upon 1h, 6h and 24h of FSC and EBSS withdrawal or rapamycin treatment (n=50-60 neurons/timepoint form n=4 biological replicates). Analysis was performed by calculating intrasynaptic over extrasynaptic LC3 levels. Scale bars: 5µm, Data represent mean ± SEM. *p < 0.05, **p < 0.005; statistical analysis by two-way ANOVA (B,E,G) or one-way ANOVA (I, K,N) with two-stage linear step-up procedure of Benjamini, Krieger and Yekutieli post hoc test.

We next examined whether nutrient deprivation induces autophagy in the adult brain and whether synapses respond differently from somatic compartments. We measured LC3-II accumulation in cytosolic and synaptosomal fractions from acute cortical slices (see Fig. S1C-E) subjected to 6h of serum withdrawal (with or without additional amino acid deprivation), glucose starvation, or rapamycin. In the presence of BafA, serum withdrawal robustly increased LC3-II levels in the cytosolic fraction (**Fig. 2D,E)** and enhanced autophagic flux (**Fig. S2C**), confirming that autophagy in non-synaptic compartments is activated under these conditions. LC3-II accumulation was also detectable in synaptosome-enriched fractions following serum withdrawal (**Fig. 2F,G, Fig. S2D**) but the effect was less pronounced than in the cytosol. In contrast, amino acid or glucose withdrawal and rapamycin had minimal effects. These data suggest that autophagy in the adult brain is most strongly triggered by serum withdrawal, and that synaptic autophagy is either less pronounced or proceeds with faster kinetics, rendering it less accessible to steady-state biochemical assays.

The difference in autophagy induction between cytosolic fraction and synapses may reflect temporal offsets, differential signaling thresholds, or faster autophagosome maturation at synapses. To dissect these dynamics, we first quantified LC3 puncta in neuronal soma after 1, 6, and 24 h of serum starvation using immunocytochemistry. LC3 puncta increased rapidly at 1 h, declined at 6 h, and become upregulated again at 24 h (**Fig. 2H, I**). To test whether this decline between 1h and 6h reflected autophagosome maturation and degradation, we used a tandem mCherry-GFP-LC3 reporter to distinguish between autophagosomes (GFP^+^/mCherry^+^) and acidified autolysosomes (GFP^−^/mCherry^+^) (**Fig. 2J, Fig. S2E**). At 2h of serum deprivation, autophagosome numbers were significantly increased (**Fig. 2K**), while autolysosome numbers were significantly reduced with both serum starvation and rapamycin (**Fig. 2L**), suggesting efficient autophagosome clearance and rapid lysosomal fusion. We next asked whether similar dynamics apply to synaptic compartments. Immunostaining revealed that LC3 levels at synaptic sites increased significantly after 6 h of serum withdrawal (**Fig. 2M, N**), with a trend already evident at 1 h. In contrast, EBSS and rapamycin had no significant effect on synaptic LC3 distribution (**Fig. S2F, G**), in line with their milder effect on the synaptic proteome (**Fig. 1**).

Together, these results demonstrate that multiple forms of metabolic stress rapidly and potently induce neuronal autophagy, with serum withdrawal being the most robust stimulus. Autophagy is activated within 1–2 h and occurs both in the soma and at synaptic terminals, albeit with distinct temporal profiles. Our data suggest that autophagy at synapses is not only temporally distinct from somatic responses, but may also be governed by specialized regulatory mechanisms.

### Acute nutrient deprivation triggers rapid recruitment and turnover of autophagic organelles at synaptic terminals

The time course of LC3 lipidation established above (Fig. 2) suggested that autophagic structures may be rapidly formed at synaptic compartments within the first 1-2 hours of serum withdrawal. To test this, we performed ultrastructural analysis of presynaptic terminals by transmission electron microscopy (TEM) after 2-hour treatments with serum-free medium and compared them to two additional metabolic stressors-EBSS, or rapamycin (**Fig. 3A**). Quantification of autophagic structures confirmed a significant increase in the number of autophagosomes at presynaptic terminals following serum deprivation (**Fig. 3B**), confirming that even short-term nutrient withdrawal is sufficient to trigger local induction of autophagy. EBSS produced a mild, non-significant increase in multivesicular bodies (MVBs), whereas rapamycin had no detectable effect, indicating that serum withdrawal is the most potent inducer of presynaptic autophagic structures in this timeframe.

**Figure 3.**
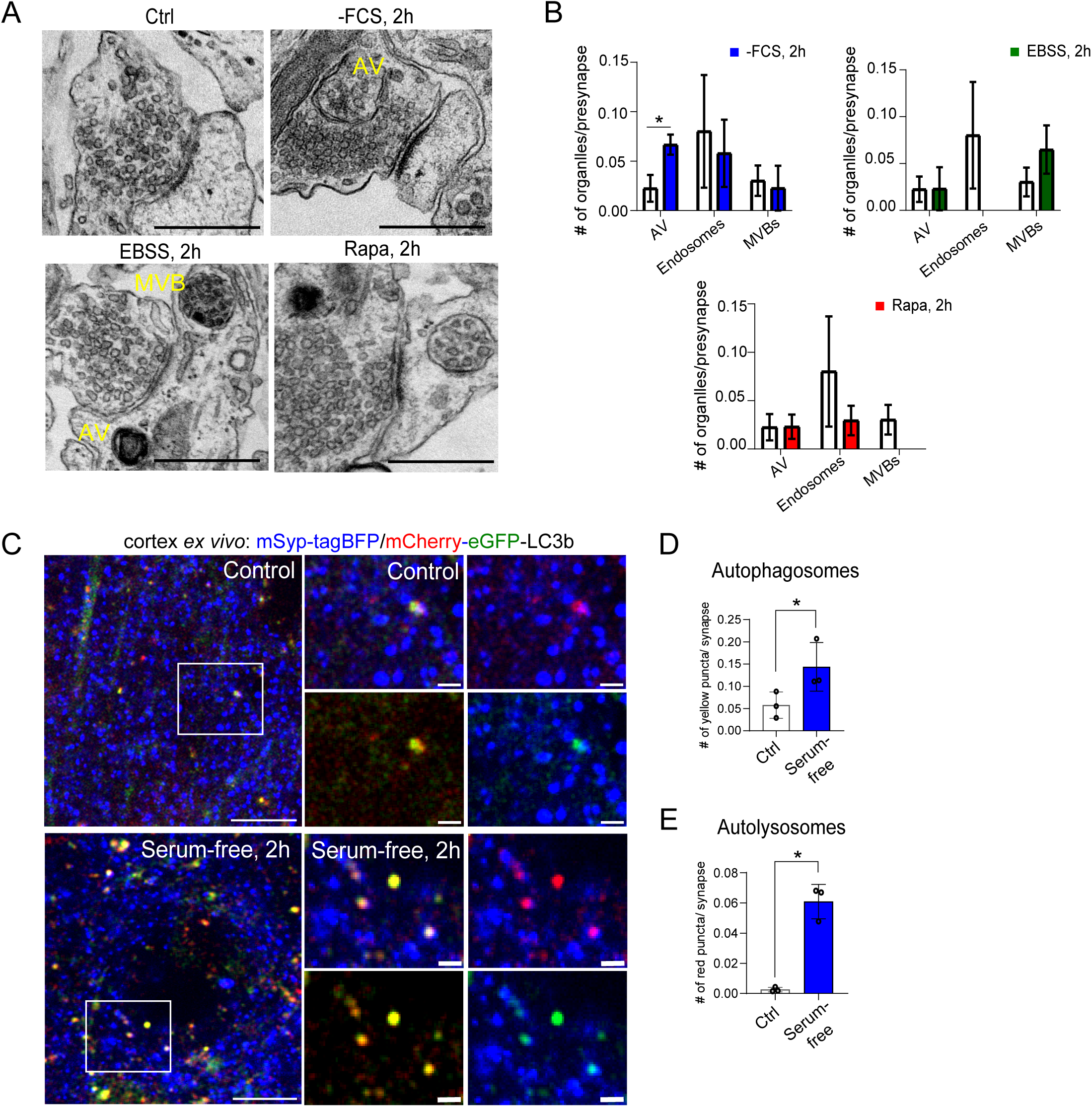
Acute nutrient deprivation triggers synaptic autophagosome formation *in vitro* and *ex vivo*. **(A)** Transmission electron micrographs of presynaptic terminals in primary cortical neurons kept either in control maintenance medium or treated for 2 h with serum-free medium, EBSS, or rapamycin (10nM). AV-autophagic structures, MVB-multivesicular bodies. Scale bars: 500 nm. **(B)** Quantification of AV, endosomes and/or MVBs in presynaptic terminals of neurons upon 2h of nutrient stress (n =74–90 terminals/condition from n=3 biological replicates). **(C)** Live imaging of cortical organotypic slice cultures transduced with mCherry-eGFP-LC3b and Syp-tagBFP (both AAV9) and kept in control maintenance medium or in the serum-free medium lacking for 2h. Scale bars: 10 µm, inserts 2µm. (**D–E**) Quantification of yellow (autophagosomes) and red-only (autolysosomes) LC3 puncta at synaptophysin-tagBFP-marked synapses (n=856 cells, 320 control cells, 536 serum starvation cells, n =6 slices from N=3 biological replicates). Data represent mean ± SEM. *p < 0.05; statistical analysis by two-way ANOVA with two-stage linear step-up procedure of Benjamini, Krieger and Yekutieli post hoc test (B) or by unpaired Student’s t-test (D).

To functionally validate these observations in living neurons, we next used live-cell imaging of cortical organotypic slice cultures co-expressing mCherry-eGFP-LC3b and the presynaptic marker synaptophysin tagged with BFP (Syp-tagBFP) (**Fig. 3C**). Serum withdrawal for 2 hours significantly increased the number of both yellow (autophagosomes) and red-only (autolysosomes) LC3 puncta at synaptic sites (**Fig. 3D,E**), suggesting enhanced autophagosome formation and rapid progression into autolysosomes. Together, these results show that in 2 hours, serum starvation can rapidly activate synaptic autophagy. The early recruitment of autophagic structures—detected both morphologically by TEM and functionally by live imaging—suggests a highly responsive and spatially targeted degradation pathway at presynaptic terminals during metabolic stress.

### ATG5-dependent autophagy contributes to synaptic proteome remodeling during nutrient stress

Having established that nutrient deprivation induces autophagy in synaptic compartments, we next sought to determine whether this process directly contributes to the remodeling of the synaptic proteome. To this end, we required a system in which autophagy could be efficiently ablated in synapses. Because constitutive *Atg5* KO mice are perinatally lethal [27], and conditional *Atg5* deletion in adult brain tissue would yield recombination only in specific synaptosomal fractions, we used cultured cortical neurons from ethanol (vehicle control)-treated (WT) and/or tamoxifen-treated *Atg5*flox:CAG-Cre^Tmx^ mice (ATG5 cKO). We chose *Atg5* because it is essential for LC3 conjugation and autophagosome formation [28], and its deletion robustly blocks macroautophagy [29, 30]. This in vitro approach enabled homogeneous and complete knockout of ATG5 (**Fig. S3A,B**), allowing reliable proteomic analysis of autophagy-dependent remodeling under nutrient stress.

To determine whether serum deprivation–induced remodeling of the synaptic proteome (see Fig. 1) depends on autophagy, we performed comparative mass spectrometry (MS) analysis of synaptosomes isolated from WT and ATG5 cKO neurons under basal and nutrient-deprived conditions (6 h serum/amino acid withdrawal). Effective deletion of ATG5 was confirmed by significantly reduced ATG5 abundance and increased p62 levels in KO synaptosomes (**Fig. S3C,D**), validating autophagy impairment. In addition, several classical autophagy cargo proteins, including TEX264, NBR1, and SEC62, accumulated in ATG5 cKO synaptosomes (**Fig. 4A**), validating autophagy impairment. Note that ATG5 values were not zero due to missing value imputation in label-free quantification workflow.

**Figure 4.**
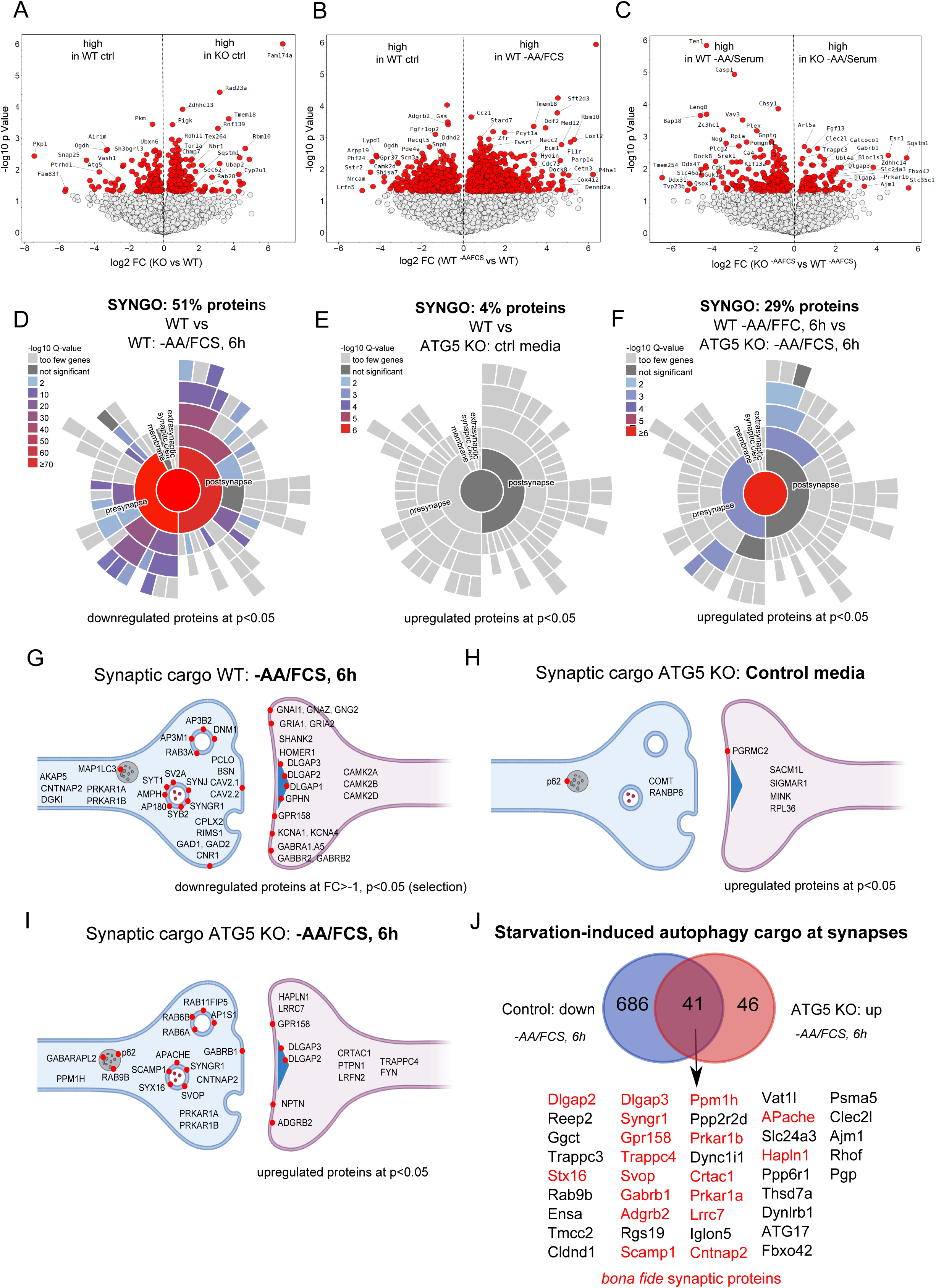
ATG5-dependent autophagy mediates nutrient-induced synaptic proteome remodeling. **(A)** Volcano plot depicting differentially expressed proteins in synaptosomes isolated from ATG5 cKO neurons maintained under basal (nutrient-rich) conditions, relative to WT controls. **(B)** Volcano plot of synaptosomes from WT neurons subjected to serum and amino acid deprivation for 6h, compared to nutrient-rich controls. **(C)** Volcano plot comparing synaptosomes from ATG5 cKO neurons and WT neurons, both maintained under nutrient-deprived conditions for 6h. The blunted proteomic response in ATG5-deficient neurons (compare to Fig. 3B) reveals impaired nutrient-induced synaptic remodeling in the absence of macroautophagy. (**D–F**) Heatmaps of SynGO-annotated synaptic proteins showing the impact of nutrient stress in WT (D) and ATG5 cKO (E,F) neurons. (**G-I**) GO enrichment analysis of starvation-responsive synaptic proteins in WT (G) and ATG5 cKO neurons under control (H) and starvation (I) conditions. In WT synapses, starvation-sensitive proteins are significantly enriched in pre- and postsynaptic compartments; this enrichment is attenuated in ATG5-deficient neurons. (**J**) Starvation-induced candidate autophagy substrates: proteins significantly (p<0.05) downregulated in WT synapses upon nutrient deprivation but significantly upregulated in ATG5 cKO neurons under the same conditions. Representative examples include PRKAR1B, SYNGR1, and DLGAP2/3. Data from n=3–4 biological replicates per condition. In (A-C) selected proteins are annotated; significance is shown in red and reflects Student’s t-test at p<0-05.

Under control conditions, WT and ATG5 KO synapses displayed only minor proteomic differences (**Fig. 4A, Table S2**). In contrast, serum deprivation triggered robust remodeling in WT synaptosomes, with widespread up- and downregulation of proteins (**Fig. 4B**). These starvation-induced changes were markedly attenuated in ATG5 KO synaptosomes (**Fig. 4C**), demonstrating that starvation-induced remodeling of the synaptic proteome requires functional autophagy. Approximately 50% of the proteins significantly downregulated in WT upon starvation were annotated as synaptic by the SynGO database [31] (**Fig. 4D**), underscoring the nutrient sensitivity of synaptic compartments. In contrast, ATG5 KO synaptosomes under control conditions showed no significant enrichment for regulated synaptic proteins (**Fig. 4E**), suggesting that basal synaptic proteostasis is largely maintained without ATG5.

Upon starvation, however, ATG5 KO synapses failed to downregulate many of the same proteins as WT (**Fig. 4F**), confirming that autophagy is required for their degradation. Functional enrichment analysis of the starvation-responsive proteome in WT synapses revealed significant representation of both pre- and postsynaptic proteins, including key cytoskeletal, scaffold, and vesicular trafficking components (e.g., GPR158, SCAMP1, SYT1, SV2A, RIMS1, CNTNAP2, GRIA1) (**Fig. 4G**). Among the most prominently regulated were proteins previously identified by us and others as autophagy substrates and regulators at synapses—such as the PKA regulatory subunits PRKAR1A and PRKAR1B [30, 32], SYNJ [33], BSN [34] and DLGAP2 [35]. These proteins were robustly downregulated in WT synaptosomes upon starvation but remained unchanged or were even upregulated in ATG5-deficient synapses upon nutrient stress (**Fig. 4H-J**), suggesting their degradation is mediated by autophagy. Together, these results support ATG5-dependent autophagy is a principal contributor to nutrient-induced synaptic proteome remodeling.

### Serum withdrawal promotes dynamic recruitment of the LC3 lipidation machinery to synapses

To dissect how the autophagy machinery is mobilized at synapses, we investigated the distribution and dynamics of the LC3 lipidation complex—comprising ATG5, ATG12, and ATG16L1—in response to metabolic stress. We first performed immunoprecipitation of endogenous ATG12 from cytosolic and synaptosomal fractions of adult cortical slices under basal conditions (**Fig. S4A**). We found that the full ATG5–ATG12–ATG16L1 complex was readily detectable in the cytosol, whereas synaptosomes contained only the ATG5∼ATG12 conjugate but little to no ATG16L1 comparing to the cytosolic fraction (**Fig. 5A, Fig. S4A**), indicating limited steady-state presence of the complete LC3 lipidation machinery and/or exclusion of ATG16L1 from synapses at baseline. Indeed, direct immunoblot analysis confirmed that ATG16L1 levels were significantly low at synaptosomes compared to the cytosolic fraction (**Fig. 5B,C**). ATG5∼ATG12 was also significantly reduced at synapses, although still detectable (**Fig. 5C**). These findings indicate that the LC3 lipidation machinery is present at low abundance at synaptic sites under baseline conditions.

**Figure 5.**
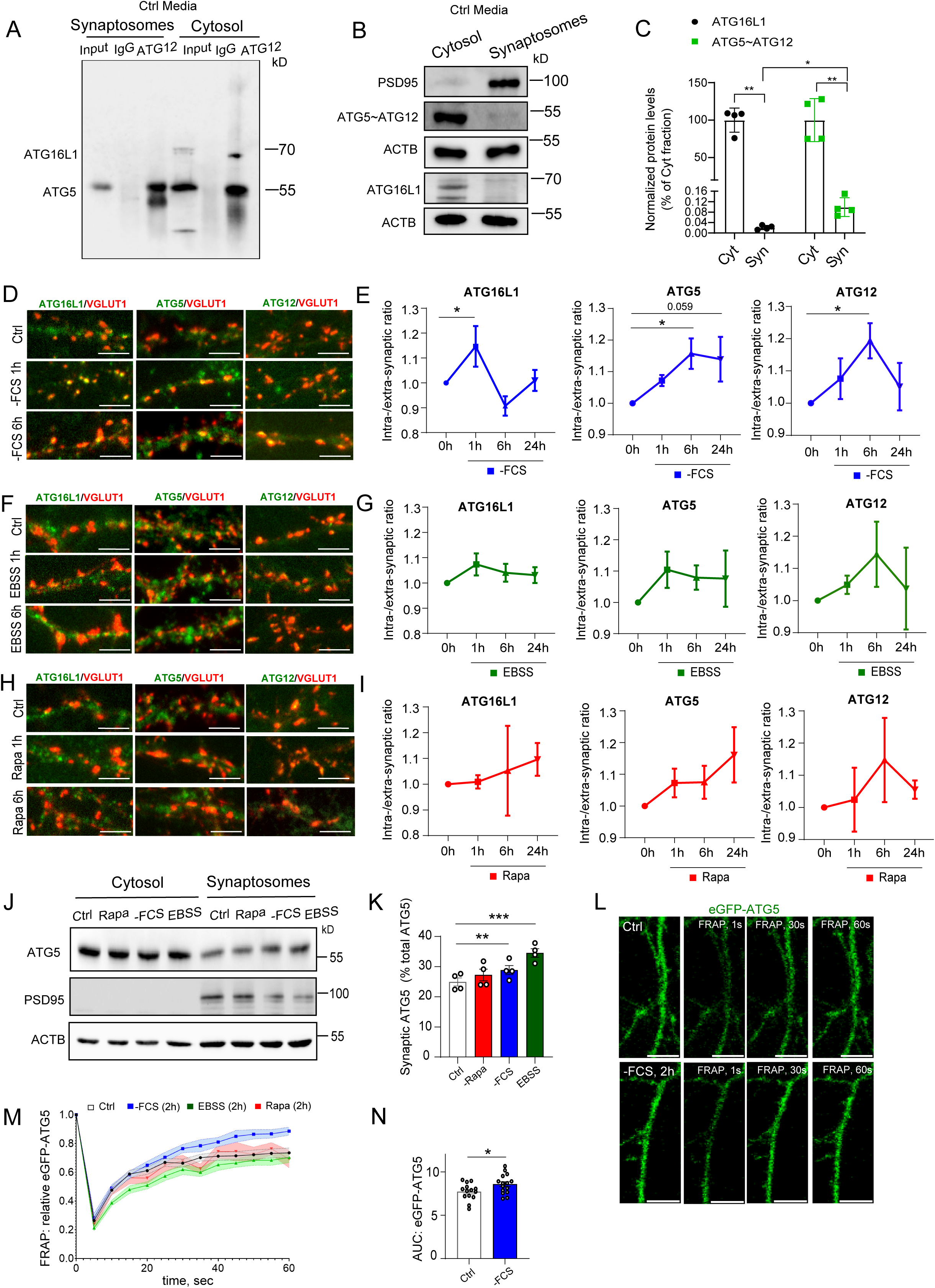
Serum withdrawal drives recruitment of the LC3 lipidation complex to synapses. **(A)** Co-immunoprecipitation of ATG12 from cytosolic and synaptosomal fractions of acute cortical slices reveals robust presence of the full ATG5–ATG12–ATG16L1 complex in the cytosol, while synaptosomes contain only the ATG5∼ATG12 conjugate with minimal detectable ATG16L1. (**B, C**) Direct immunoblotting (B) and protein quantification (C) of ATG5 and ATG16L1 from synaptosomal and cytosolic fractions confirms low steady-state abundance of ATG16L1 and reduced—but detectable—levels of ATG5 at synaptic sites. Protein levels were normalized to ACTB. n = 4 biological replicates. Baseline levels in cytosol are normalized to 100%. (**D–I**) Time course of ATG16L1, ATG5, and ATG12 recruitment to VGLUT1-marked to synaptic sites in primary cortical neurons subjected to nutrient stress. Immunofluorescence analysis was performed after 1h, 6h, and 24h of serum withdrawal (D, E), EBSS treatment (F, G), or rapamycin (H, I). Synaptic recruitment was quantified by calculating the ratio of intrasynaptic to extrasynaptic fluorescence intensities for each protein using VGLUT1 as a synaptic marker. Data represent 40–60 neurons per time point from n=3–5 independent biological replicates. Scale bars: 5µm. (**J,K**) Immunoblotting (J) and protein quantification (K) of ATG5 in synaptosomal and cytosolic fractions from cortical slices treated for 6h with serum-free medium (-FCS), EBSS or rapamycin (10 nM) or kept in control medium. Protein levels were normalized to ACTB. n=4 biological replicates. (**L–N**) FRAP (fluorescence recovery after photobleaching) analysis of eGFP-tagged ATG5 in live primary cortical neurons showing increased mobility of ATG5 upon serum deprivation. Representative images (L) show ATG5 puncta before and after bleaching under control and – serum deprivation (2h) condition. Time-lapse imaging was performed at 1s, 30s, and 60s post-bleaching. Quantification of fluorescence recovery (M) shows faster recovery in serum-starved neurons, indicative of ATG5 increase trafficking. Area under the curve (AUC) of fluorescence recovery is significantly increased after serum withdrawal (N). Data from n = 3 independent biological replicates. Scale bars: 10 µm. Data represent mean ± SEM. *p < 0.05, **p < 0.01, ***p < 0.001; statistical analysis by two-way ANOVA (C) or one-way ANOVA with Holm–Sidak post hoc test (E, G, I, K, N), or by unpaired Student’s t-test (N).

We next tested whether nutrient deprivation alters this distribution. Serum withdrawal induced a time-dependent recruitment of ATG16L1, ATG5, and ATG12 to synapses, with intra-/extrasynaptic ratios peaking at 1–6 hours and returning toward baseline by 24 hours (**Fig. 5D,E**). EBSS produced a similar trend, but the changes did not reach statistical significance (**Fig. 5F,G**). In contrast, rapamycin did not significantly alter the synaptic localization of any LC3 lipidation component (**Fig. 5H,I**), consistent with its limited effect on synaptic autophagy observed above (**Figs. 1–3**). Quantitative immunoblotting of synaptosomal and cytosolic fractions after 6 h of treatment confirmed this recruitment: both serum deprivation and EBSS increased synaptic ATG5 levels relative to cytosolic levels, whereas rapamycin had no effect (**Fig. 5J,K**).

Finally, to assess the mobility and stabilization of ATG5 upon serum starvation, we performed fluorescence recovery after photobleaching (FRAP) analysis in cortical neurons expressing eGFP–ATG5 (**Fig. 5L**). Under control conditions, eGFP–ATG5 displayed rapid recovery, indicating high turnover. Upon serum withdrawal, this recovery was significantly increased (**Fig. 5M,N**), suggesting that nutrient stress enhances ATG5 mobility in neurons.

Together, these findings demonstrate that serum deprivation triggers a temporally coordinated recruitment of the LC3 lipidation machinery at synapses. The near-absence of ATG16L1 in synapses at baseline, and its inducible enrichment during stress, suggests that synaptic autophagy initiation is tightly regulated and likely requires active trafficking of essential components into the compartment.

### Autophagy machinery recruitment to synapses is mediated by Rab5b

To identify molecular mediators of LC3 lipidation machinery recruitment to synapses, we investigated starvation-induced interactors of ATG16L1. We performed immunoprecipitation of endogenous ATG16L1 from cytosolic and synaptosomal fractions of control and serum-deprived cortical slices, followed by label-free quantitative proteomics (**Fig. 6A**). Serum deprivation markedly re-shaped the ATG16L1 interactome, particularly in synaptosomes, where a distinct set of synaptic proteins was co-enriched with ATG16L1 upon nutrient deprivation (**Fig. 6B,C**). Among the 36 starvation-induced binding partners, we identified several proteins previously implicated in autophagy regulation, including Ubqln1/2 [36], Endophilin A1 (EndoA1) [37], ERC1/Clasp2 [29] and Rab5 [38] (**Fig. 6D**, **Extended Table S3**).

**Figure 6.**
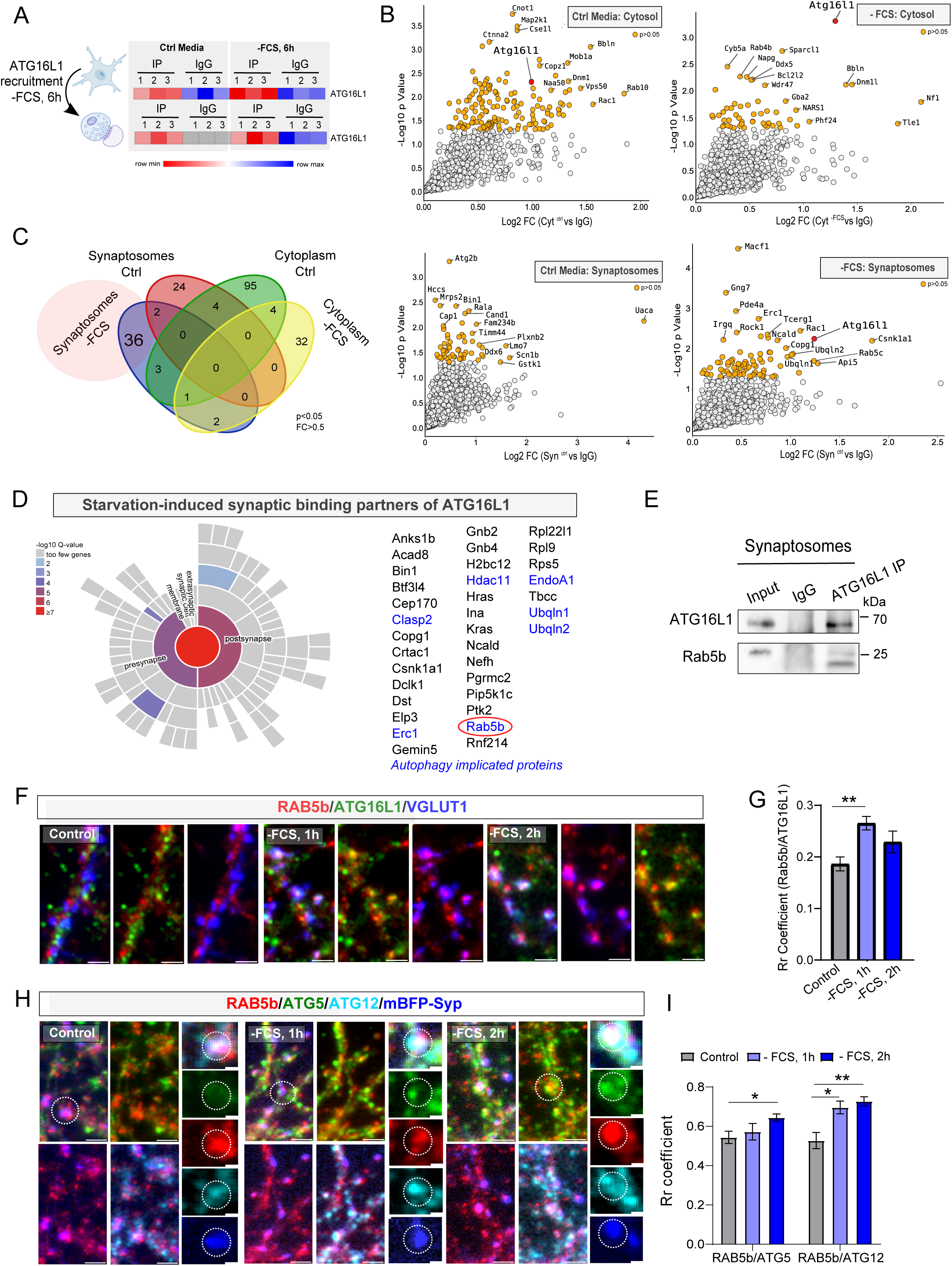
Rab5b mediates starvation-induced recruitment of the autophagy machinery to synapses. **(A)** Experimental workflow: ATG16L1 was immunoprecipitated from cytosolic and synaptosomal fractions of control and serum-starved (6h) adult cortical slices followed by LC–MS/MS–based identification of interacting proteins. **(B)** Volcano plots showing differentially enriched ATG16L1 interactors in cytosolic and synaptosomal (fractions under nutrient-rich versus starvation conditions. Serum deprivation induces a distinct set of ATG16L1 interactions in synaptic compartments. n=3 biological replicated per condition. Selected proteins are annotated; significance is shown in yellow and reflects Student’s T-test-adjusted at p<0-05. **(C)** Venn diagram depicting the overlap of steady-state and starvation-induced ATG16L1 interactors in cytosolic and synaptosomal fractions, revealing a distinct set of synapse-specific binding partners recruited under nutrient stress. **(D)** Among the top starvation-enriched ATG16L1 interactors in synaptosomes is Rab5b, a small GTPase associated with early endosomes and membrane trafficking. **(E)** Western blot validation confirms increased interaction between endogenous Rab5b and ATG16L1 in synaptosomes isolated from acute cortical slices. ATG16L1 was immunoprecipitated from synaptosomes followed by Western blotting for Rab5b. (**F-H)** Time-resolved immunocytochemistry (F,H) and quantification (G,I) of Rab5b colocalization with ATG16L1, ATG5, and ATG12 at VGLUT1-positive synapses in cultured cortical neurons during the early stages of serum deprivation (1–2h). Quantification reveals a temporally ordered recruitment of the LC3 lipidation machinery to presynaptic compartments: Rab5b–ATG16L1 colocalization peaks at 1h, followed by Rab5b–ATG5 at 2h. 18–29 neurons per timepoint from n=3 biological replicates. Scale bars, 2µm, magnified areas: 0.5µm. Data represent mean ± SEM. *p<0.05, **p<0.005; statistical analysis by two-way ANOVA (I) or one-way ANOVA with Holm–Sidak post hoc test (G).

Rab5 proteins (Rab5a, Rab5b, and Rab5c) are members of the small GTPase family involved in early endosome formation, membrane trafficking, and synapse function [39–42]. While Rab5a has been widely implicated in classical endocytic and autophagic pathways [43–46], Rab5b is less well characterized but has been reported to mediate several endocytic events [47, 48], including cargo-specific trafficking at synapses [49–51]. Given these properties, we hypothesized that Rab5b may serve as a synapse-specific trafficking adaptor for the LC3 lipidation machinery during nutrient stress. To test this, we first validated the interaction between ATG16L1 with Rab5b by Western blot and confirmed that this interaction occurs in synaptic compartments (**Fig. 6E**). To further assess the spatial and temporal dynamics of this interaction, we used immunocytochemistry in cultured cortical neurons and quantified colocalization of Rab5b with ATG16L1, ATG5, and ATG12 at synaptic sites marked by VGLUT1 and/or synaptophysin-mBFP (**Fig. 6G–J**). Notably, colocalization between Rab5b and ATG16L1 and Rab5b and ATG12 was significantly increased already at 1 h of serum deprivation, followed by enhanced Rab5b–ATG5 colocalization at 2 h, indicating a temporally ordered recruitment of LC3 lipidation components to synaptic compartments.

Together, these results suggest a model in which Rab5b-positive early endosomes act as carriers for the LC3 lipidation machinery, delivering ATG16L1–ATG12–ATG5 complexes to presynaptic terminals under nutrient stress.

### Nutrient deprivation enhances Rab5b–ATG16L1 co-trafficking and requires microtubule-based transport for synaptic recruitment

To dissect the transport dynamics of Rab5-positive endosomes during nutrient stress, we turned to live-cell imaging of primary neurons co-expressing mCherry-tagged Rab5b and mEmerald-tagged ATG16L1. Under control conditions, Rab5b vesicles exhibited modest mobility and limited co-trafficking with ATG16L1 (**Fig. 7A**). However, within 1–2 h of serum withdrawal, the velocity of Rab5b-positive vesicles doubled (**Fig. 7B**), coinciding with a pronounced increase Rab5b-ATG16L1 colocalization (**Fig. 7C**). Quantification of vesicle trajectories revealed a marked increase in co-trafficking between Rab5b and ATG16L1 after 1 h of serum withdrawal (**Fig. 7D**), indicating that Rab5b-positive endosomes actively transport ATG16L1 under metabolic stress.

**Figure 7.**
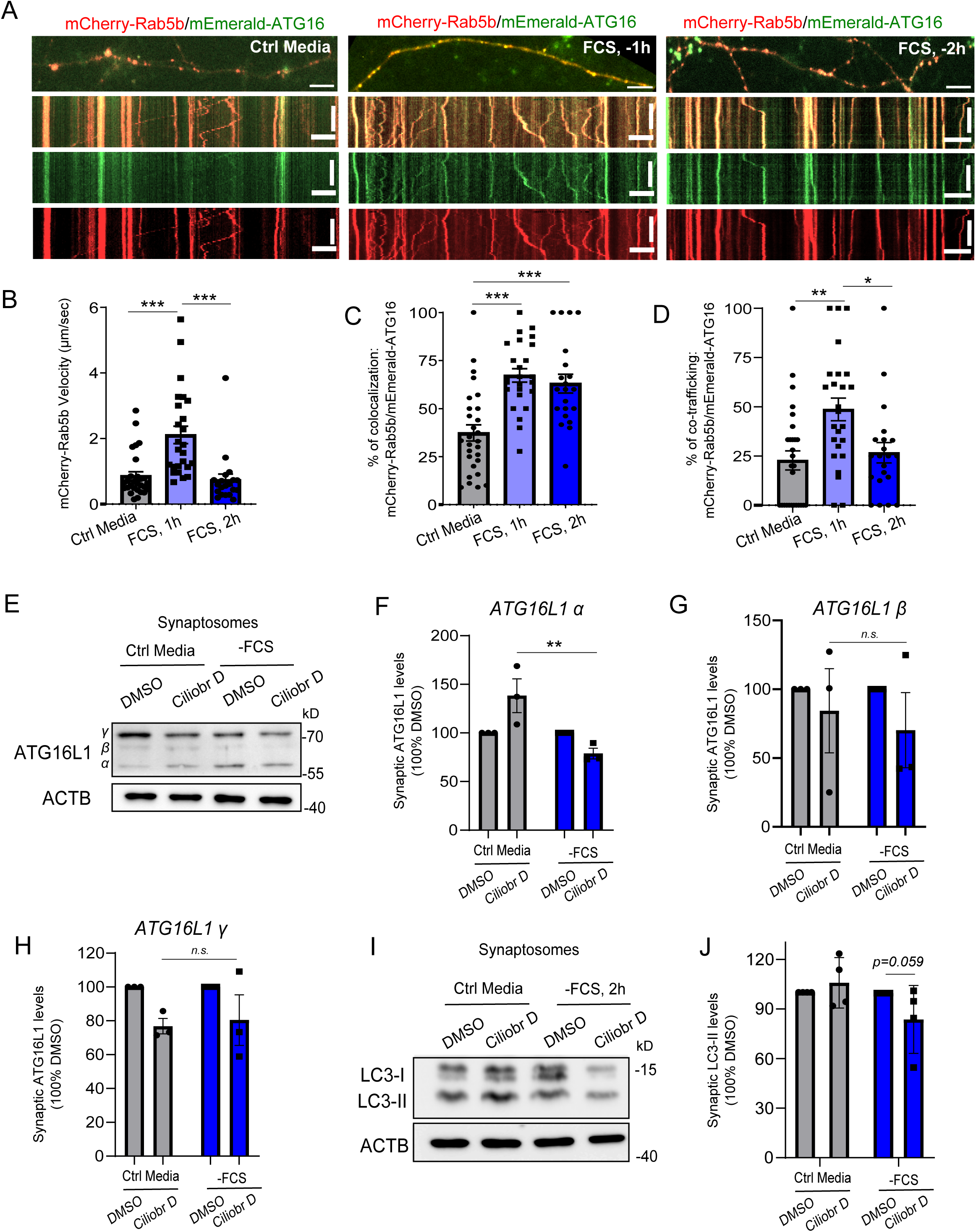
Rab5b-mediated trafficking of ATG16L1 to synapses is dynein-dependent and enhanced by nutrient stress. **(A)** Presentative live-cell images and corresponding kymographs of primary cortical neurons co-expressing mCherry-Rab5b and mEmerald-ATG16L1, maintained in control medium or subjected to serum withdrawal (–FCS) for 1–2 h. Scale bars, x µm (image) and y s (kymograph). Scale bars: 5µm (upper panels), 5µm x 20sec **(B)** Rab5b vesicle velocity in axons increases after 1 h of serum withdrawal (20–28 neurons per condition, n = 4 independent cultures). **(C)** Percentage of Rab5b puncta positive for ATG16L1 increases at 1 h and 2 h of serum withdrawal (20–28 neurons per condition, n = 4). **(D)** Co-trafficking of Rab5b–ATG16L1 vesicles increases after 1 h of serum withdrawal (20–28 neurons per condition, n = 4). **(E)** Immunoblot of synaptosomal fractions from neurons treated with 100 nM of Ciliobrevin D (Ciliobr D) and ± 6 h serum withdrawal, probed for total ATG16L1. (**F-H**) Western blot analysis of ATG16L1 isoforms (α, β, γ) in control and serum-starved synaptosomes upon 6h treatment with ciliobrevin D. Starvation-induced enrichment of the ATG16L1 α isoform at synapses was abolishes by dynein inhibition (CTRL+ Ciliobr D vs FCS+ Ciliobr D, p = 0.019). DMSO baseline controls are normalized to 100 (n = 3 independent biological replicates). **(I,J)** Immunoblot analysis (I) and quantification (J) of LC3 levels in synaptosomal fractions upon ciliobrevin D treatment with or without serum starvation. n=4 biological replicates. Data represent mean ± SEM. *p<0.05, **p<0.005; statistical analysis by two-way ANOVA (F-H, J) or one-way ANOVA with Holm–Sidak post hoc test (B-D).

Rab5-positive early endosomes are known to rely on dynein for retrograde microtubule transport in neurons [52, 53], and our data indicate that Rab5b vesicles serve as carriers for ATG16L1 during nutrient stress (**Fig. 6**). We therefore reasoned that dynein-dependent microtubule-based transport is required for recruitment of autophagy machinery to synapses. To test this, we inhibited dynein using Ciliobrevin D-a selective inhibitor of cytoplasmic dynein ATPase activity that does not affect kinesin-driven motility- and assessed ATG16L1 levels in synaptosomal fractions following serum withdrawal (**Fig. 7E**). ATG16L1 was detected as multiple isoforms in synaptosomes: a prominent band at ∼58 kDa corresponding to ATG16L1α, and a higher molecular weight species at ∼70 kDa corresponding to the ATG16L1γ variant, consistent with previous reports describing distinct ATG16L1 isoforms generated through alternative splicing [54]. The specificity of these bands was confirmed by their prominent reduction in Atg16L1 cKO and ATG5 cKO brain lysates (**Fig. S4B,C**), validating the antibody and confirming isoform dependence on intact autophagy machinery.

Ciliobrevin D treatment significantly reduced starvation-induced accumulation of ATG16L1α isoform at synapses, without affecting its levels under control (non-starved) conditions (**Fig. 7F**). This indicates that microtubule-based, dynein-dependent transport is required for the synaptic delivery of ATG16L1 during nutrient stress. In line with this, LC3-II levels at synapses trended downward after Ciliobrevin D treatment during 2h of serum withdrawal (**Fig. 7I,J**), although the effect did not reach statistical significance (p = 0.059). Similar modest reductions were observed in independent replicates after 6 h starvation (**Fig. S4D,E**). These limited effects on LC3-II accumulation likely reflect the highly dynamic nature of autophagosomes at synapses. Specifically, while ciliobrevin D inhibits the dynein-dependent delivery of ATG16L1 via Rab5b-positive endosomes, other trafficking routes, such as the kinesin-driven, RAB39-mediated transport of ATG9 from soma to synapse [55], may remain unaffected. Additionally, mature autophagosomes formed prior to dynein inhibition could still be cleared, further attenuating the net biochemical signal in LC3-II levels.

Taken together, our data suggest that Rab5b-positive endosomes retrogradely transport pre-autophagic machinery from distal axonal segments to synapses. Nutrient stress enhances the velocity and co-trafficking of Rab5b and ATG16L1, enabling targeted delivery of autophagosome precursors to metabolically challenged synaptic compartments. The selective regulation of the 58 kDa ATG16L1 isoform further suggests isoform-specific stabilization or transport during early autophagy induction.

### Nutrient deprivation modulates neuronal activity and enhances synaptic remodeling in vivo

Given our findings that nutrient deprivation induces synaptic autophagy and proteome remodeling (**Figs. 1–7**), we next asked whether these molecular changes translate into altered neuronal function. Because synaptic composition critically shapes excitability and neurotransmission [56, 57], we hypothesized that nutrient stress might modulate neuronal activity. To test this, we performed calcium imaging in primary neurons transduced with AAV9-hSyn-GCaMP7f (**Fig. 8A**) and subjected to acute metabolic stress (2 h) using either serum-free medium, EBSS, or rapamycin, followed by high-frequency stimulation (4×1sec pulses at 100Hz). Serum withdrawal led to a prominent reduction in stimulus-evoked GCaMP7f transients, while EBSS and rapamycin caused milder but detectable decreases (**Fig. 8B**). Quantification of the area under the curve (AUC) confirmed a significant reduction in calcium responses following serum deprivation (**Fig. 8C**), indicating that short-term nutrient stress dampens neuronal excitability, possibly via autophagy-dependent synaptic remodeling.

**Figure 8.**
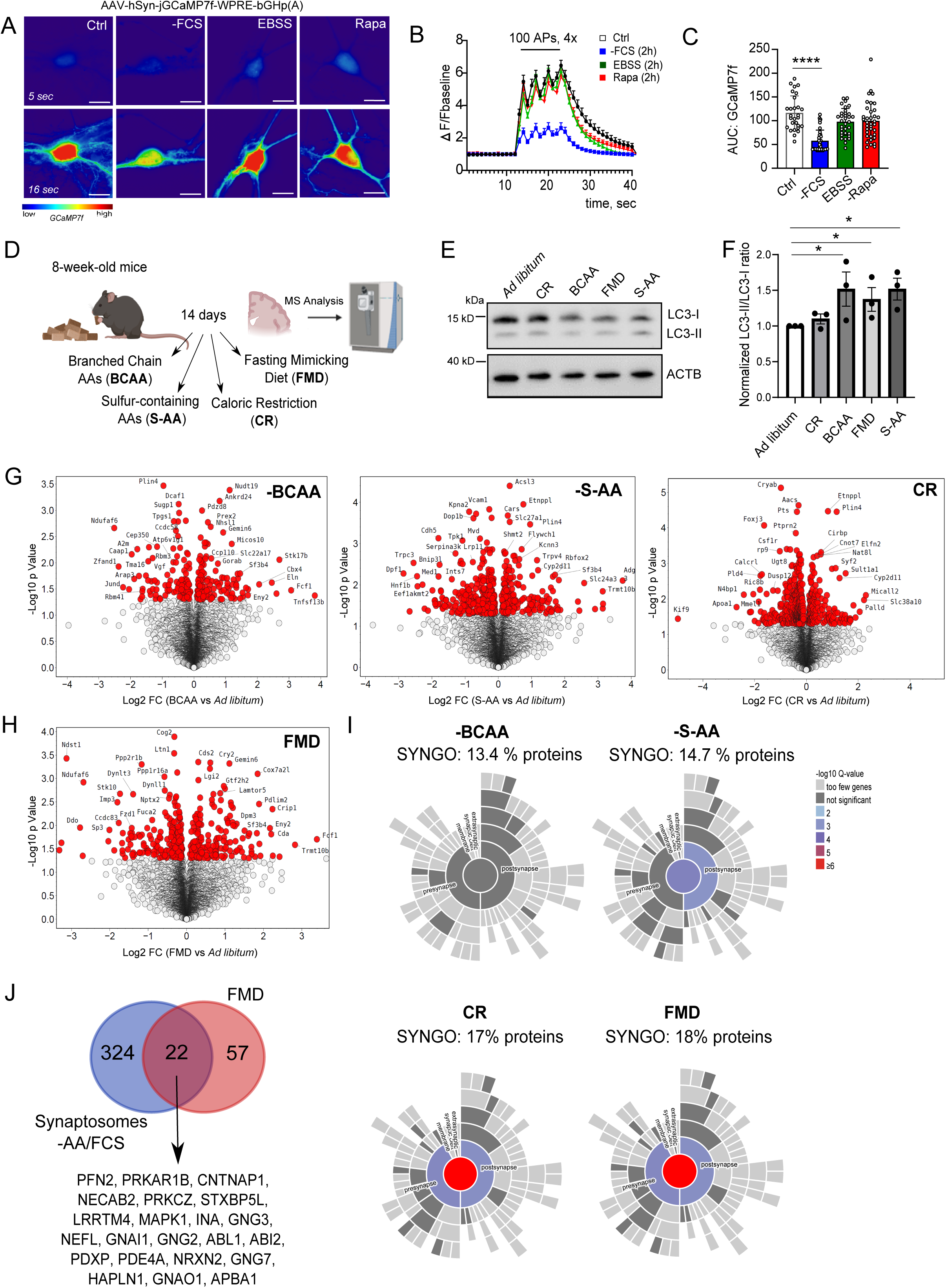
Nutrient restriction reshapes neuronal activity and autophagy state *in vivo*. (**A-C)** GCaMP7f-based Ca^2+^ imaging performed in primary cortical neurons transduced with hSyn[jGCaMP7f. neurons were subjected to 2h using serum-free medium (–FCS), EBSS, or rapamycin (10 nM) prior to high-frequency stimulation (4×1 s trains at 100 Hz). Representative heatmaps of calcium responses in individual neurons and baseline-normalized GCaMP7f fluorescence (ΔF/F_0_) under each condition are shown in (A) and (B) respectively. Quantification of area under the curve (AUC) reveals significant reduction in calcium responses upon serum withdrawal (C). Data from >28 neurons per group pooled from N=2 independent cultures. Scale bar: 5µm. **(D)** Schematic overview of in vivo dietary paradigms and proteomics workflow. Eight-week-old mice were subjected for 14 days to caloric restriction (CR), fasting-mimicking diet (FMD), or amino acid–restricted diets lacking branched-chain amino acids (BCAA) or sulfur-containing amino acids (S-AA). Cortices were harvested for biochemical and proteomic analyses (see also Suppl. Table S5). **(E,F)** Representative immunoblots showing LC3-I and LC3-II levels in cortical lysates across diet groups. Quantification of LC3-II/LC3-I ratios normalized to *ad libitum* controls reveals significant increases under FMD, BCAA, and S-AA conditions. n = 3 biological replicates. **(G,H)** Volcano plots showing differentially expressed proteins in cortex across each diet group (BCAA, S-AA, CR, FMD) versus *ad libitum*-fed controls. Selected proteins are annotated; significant hits highlighted in red. n=4 biological replicates. **(I)** Heatmaps display expression changes of SynGO-annotated synaptic proteins (pre-, post-, and extrasynaptic) across all diets. FMD and CR induce pronounced synaptic changes, while BCAA and S-AA exert limited effects. Percentages of regulated proteins with synaptic annotations are indicated for each condition. **(J)** Overlap between in vitro starvation-sensitive synaptic autophagy cargo and in vivo fasting targets. Venn diagram and list of 22 proteins significantly downregulated both in WT neurons upon 6h starvation (see Fig. 4J) and in cortical lysates of FMD-fed mice. Data are mean ± SEM; ****p<0.00001; statistical analysis by one-way ANOVA with Holm–Sidak post hoc test.

Finally, we asked whether nutrient restriction impacts the brain proteome *in vivo*. We used (i) a fasting-mimicking diet (FMD) designed to trigger coordinated nutrient deprivation and autophagy induction, (ii) caloric restriction (CR) as a broadly applied metabolic stressor, and (iii) amino acid–restricted diets either restricted in branched-chain amino acids (BCAA) or fully lacking sulfur-containing amino acids (S-AA) to inhibit mTORC1, a key negative regulator of autophagy [58–64]. Adult mice were subjected for 14 days to various dietary regimens for two weeks in a randomized manner (**Fig. 8D, Extended Table S4**). Since Baf A was not administered *in vivo* in these mice, we assessed LC3-II/LC3-I ratios as a proxy for autophagy flux. Among the different diets, FMD, BCAA, and S-AA restriction induced significant flux changes in LC3-II/LC3-I ratio in cortical lysates (**Fig. 8E,F**), suggesting enhanced autophagic activity. Proteomic analysis revealed that FMD—and to a lesser extent CR—induced pronounced remodeling of the synaptic proteome (**Fig. 8G-I**). Of the significantly deregulated proteins in CR- and FMD-treated mice, 17% (95/558) and 18% (79/440), respectively, mapped to synaptic compartments (**Fig. 8I**). In contrast, mTORC1-inhibiting diets such as a BCAA-restricted diet had minimal impact on synaptic composition, suggesting that mTORC1-dependent autophagy may play cargo-specific roles outside synaptic compartments, and that additional nutrient cues beyond mTOR inhibition are required to engage synaptic autophagy.

FMD altered a broad array of pre- and postsynaptic proteins, including those involved in vesicle cycling, receptor trafficking, and plasticity (**Fig. S5A**), consistent with targeted proteostatic remodeling. Importantly, 22 synaptic proteins differentially expressed in FMD-treated animals overlapped with starvation-induced autophagy cargo identified in Fig. 4J. Among these, we identified the regulatory subunit 1B of the PKA complex (PRKAR1B), which was previously identified by us as a synaptic autophagy substrate during nutrient deprivation [30], providing further *in vivo* validation of autophagy-dependent synaptic remodeling. Despite robust synaptic proteome remodeling, chronic dietary restriction did not significantly alter pS6 (Ser235/236) levels in cortex (**Fig. S5B,C**), consistent with prior reports that mTORC1 activity rebounds or stabilizes during prolonged nutrient restriction [65, 66].

Taken together, these results demonstrate that acute nutrient deprivation reduces neuronal excitability, whereas chronic dietary restriction *in vivo* triggers synapse-enriched changes within cortical proteomes that overlap with starvation-induced autophagy cargo. While future work is needed to validate the synaptosomal specificity of these changes and their dependence on neuronal autophagy *in vivo*, our findings nonetheless highlight a physiologically relevant mechanism by which nutrient cues remodel synaptic composition and function via compartmentalized autophagy.

## Discussion

Synaptic proteostasis is essential for maintaining neuronal integrity and function, particularly in response to metabolic fluctuations [67]. Here, we identify nutrient stress as a potent regulator of synaptic composition and function, acting through rapid, autophagy-dependent remodeling of the synaptic proteome. By systematically dissecting the temporal and spatial dynamics of autophagy induction in neurons, we reveal that synapses are both sensitive to nutrient availability and capable of mounting a rapid autophagic response to maintain proteome integrity.

mTORC1 is a well-established negative regulator of autophagy, operating primarily through phosphorylation of ULK1, which prevents its activation by AMPK and blocks autophagosome formation [22]. While pharmacological or nutrient-based mTORC1 inhibition robustly induces autophagy in many cell types [21, 68–74], its role in neurons—especially at synapses—remains more ambiguous. Studies have shown that rapamycin can induce autophagic vacuoles in various neuronal populations [75–80] and has beneficial effects in neurodegeneration models [81]. Yet, others report inconsistent or modest autophagy responses [26, 82]. Our data shed light on this issue by comparing the effects of nutrient withdrawal and mTORC1 inhibition on synaptic proteostasis. Mass spectrometry–based profiling revealed that nutrient deprivation, particularly serum withdrawal, causes robust remodeling of the synaptic proteome (Fig. 1), whereas rapamycin-induced mTORC1 inhibition produces a comparatively limited effect. In vivo, 2-week dietary restriction of BCAAs—an established mTORC1-inhibiting condition [83]-—did not significantly affect synaptic composition, despite upregulation of autophagy flux. In contrast, FMD and CR both robustly remodeled the synaptic proteome. These results suggest that while BCAA withdrawal induces autophagy, the cargo of resulting autophagosomes may largely exclude synaptic proteins. Thus, mTORC1-regulated autophagy appears to exert cargo-selective functions, and synaptic responses to metabolic stress may rely on additional nutrient-sensing pathways beyond mTORC1. We acknowledge, however, that while our findings indicate limited modulation of synaptic proteome under rapamycin, we did not directly assay mTORC1 target engagement at synapses. Moreover, alternative nutrient-sensing mechanisms, such as the AMPK–ULK1 axis [22], may also contribute to the induction of synaptic autophagy.

We show that nutrient deprivation induces autophagy both globally and at synapses (Figs. 2,3). Serum withdrawal rapidly increased LC3 lipidation and autophagosome trafficking, with synaptic autophagy peaking within 1–2 hours. This is consistent with prior studies showing rapid autophagic responses at presynaptic terminals following localized damage or starvation [79, 84], and the ordered recruitment of core autophagy factors (ATG13, ATG5, LC3) within minutes in neurons and other cells [85, 86]. These findings indicate that synaptic autophagy is rapidly and locally regulated, in line with evidence of compartmentalized autophagic flux in neurons [26].

Mechanistically, we find that nutrient stress drives the spatiotemporally regulated recruitment of the LC3 lipidation complex (ATG5–ATG12–ATG16L1) to synapses (Fig. 5). This process is orchestrated by Rab5b-positive early endosomes, which serve as carriers of autophagy machinery under starvation (Figs. 6–7). Rab5b-positive endosomes are canonically associated with early endosome trafficking and exhibit dynein-biased bidirectional transport in neurons [52, 53, 87]. While dynein is typically linked to retrograde trafficking of mature autophagosomes [88], our data suggest that Rab5b vesicles also use dynein-dependent transport to deliver pre-autophagosome machinery—including the LC3 lipidation complex—to synaptic compartments. This supports a model in which autophagy precursors are retrogradely trafficked from distal axonal regions to presynaptic terminals via Rab5b-positive early endosome carriers.

Prior work has demonstrated that ATG16L1-positive vesicles can associate with the AP-2/clathrin at the plasma membrane thereby contributing plasma membrane-derived material to early autophagosomal precursors [89]. Our data suggest that serum withdrawal may promote endocytosis at presynaptic terminals, thereby enhancing ATG16L1 recruitment to Rab5b-positive early endosomes during nutrient stress. Consistent with this, rapid starvation-induced autophagy substrates have been shown to associate with Rab5-positive endosomes in non-neuronal cells [90]. Moreover, endosome-mediated mitochondrial clearance has been reported to initiate upstream of autophagy via Beclin1-dependent Rab5 activation [91]. While it remains unclear whether similar pathways operate at synapses, our findings are consistent with previously described mechanisms of amphisome formation at synapses, in which endosomes interface with presynapse–originating autophagosomes to regulate synaptic signaling [92, 93].

Interestingly, we observe preferential role for the ATG16L1α isoform in Rab5b-mediated recruitment to synapses, suggesting isoform-specific functional specialization. Prior studies showed that the ATG16L1β isoform is dispensable for starvation-induced LC3 lipidation but plays a role in LC3 conjugation under perturbed endosomal conditions. In contrast, ATG16L1α, which lacks the C-terminal membrane-binding region present in β, supports canonical autophagy via its N-terminal amphipathic helix [94]. Future studies are warranted to determine whether ATG16L1α selectively contributes to synaptic autophagosome formation or cargo selection under metabolic stress in neurons.

This Rab5b-dependent delivery mechanism is likely distinct from that of other autophagy-related vesicles such as those carrying ATG9 [95]. Indeed, ATG9 is absent from Rab5-positive early endosomes at steady state [96], suggesting that ATG9 and LC3 lipidation components are delivered by separate, specialized membrane carriers. In contrast to the retrograde Rab5b–ATG16L1 route, ATG9 vesicles undergo kinesin-mediated anterograde transport from the soma, a pathway which can be inhibited by Rab39 [55, 95]. These complementary trafficking routes highlight the spatial complexity of autophagy initiation at synapses and suggest that synaptic autophagy relies on the coordinated delivery of distinct precursor pools.

Additional regulatory inputs may further modulate these trafficking events. In non-neuronal cells, recruitment of the LC3 lipidation machinery is highly sensitive to PI3K inhibition: wortmannin treatment abolishes nearly all ATG5 and ATG16L1 puncta while sparing a subset of ULK1 foci [86], suggesting a PI3K-dependent mechanism of LC3 machinery assembly at synapses. Moreover, mTOR inhibition has been shown to enhance interactions between Rab11a and ATG9, promoting LC3-positive vesicle formation in response to NMDA receptor stimulation [97], further underscoring the complexity of membrane trafficking regulation during autophagy initiation in neurons. We acknowledge that serum withdrawal simultaneously removes a broad array of components—including growth factors, hormones, lipids, and carrier proteins—and that the stronger effects observed with serum-free medium compared to EBSS (Fig. 1) may reflect the loss of trophic support rather than nutrient depletion alone. Indeed, withdrawal of specific growth factors such as IGF and FGF has been shown to directly influence autophagy induction [98, 99]. However, because serum is only partially defined in terms of its composition and exact concentrations, we are currently unable to pinpoint which specific factors contribute to the observed induction of synaptic autophagy upon serum deprivation.

Functionally, ATG5-deficient neurons failed to remodel their synaptic proteome upon nutrient deprivation, confirming that macroautophagy machinery is required for this adaptation (Fig. 4). Nutrient deprivation also reduced neuronal excitability (Fig. 8A–C), further linking autophagy to functional outcomes. Prolonged fasting in vivo led to significant remodeling of synaptic proteins, including 22 proteins identified as starvation-induced autophagy cargo (Fig. 4J, Fig. 8J), among them PRKAR1B, a PKA regulatory subunit we previously validated as a synaptic autophagy substrate [30]. This provides further *in vivo* evidence that nutrient stress engages synapse-targeted autophagy. Our data also align with prior studies showing that nutrient deprivation causes selective degradation of synaptic proteins, disrupts synaptic vesicle pools, and impairs vesicle release [25]. Many of the nutrient-sensitive proteins we identified are involved in vesicle cycling, neurotransmitter release and receptor trafficking, reinforcing the idea that metabolic stress directly impacts the machinery of synaptic transmission and plasticity. While our chronic dietary restriction experiments cannot fully exclude a glial contribution to cortical autophagy flux, the consistent overlap with autophagy substrates defined in neurons, along with the preservation of synapse-enriched changes, strongly supports a neuron-autonomous synaptic response.

Collectively, these findings position autophagy as a central regulator of synaptic proteostasis under metabolic stress. The spatiotemporal recruitment of autophagy machinery, particularly via Rab5b-mediated trafficking of LC3 lipidation components, underscores the role of synaptic compartments as active sites of nutrient-sensitive degradation. This has broad implications for understanding how metabolic fluctuations shape synaptic plasticity and how defects in autophagy might contribute to synaptic dysfunction in neurodegenerative and metabolic brain disorders.

## Material and Methods

### Animals and Ethical Approval

All animal experiments were approved and performed according to the regulations of the LAVE (Landesamt für Verbraucherschutz und Ernährung, Nordrhein-Westfalen), Germany guidelines (AZ: 81-02.04.2020.A418, 81-02.05.40.20.075, 81-02.04.2021.A132, 81-02.04.2021.A067, 81-02.04.2022.A116 and 2019-A345). Mice were maintained in a pathogen-free environment in ventilated polycarbonate cages and housed in groups of five animals per cage with constant temperature and humidity at 12-h/12-h light–dark cycles. Food and water were provided *ad libitum*. Mice used in this study were WT C57BL/6J and conditional tamoxifen[inducible ATG5 KO mice (*Atg5*flox/flox: B6.Cg-Tg(CAG-Cre/Esr1*)5Amc/J) were previously described [29]. **Suppl. Table 2** indicates genotyping primers used to genotype the animals. In accordance with the SAGER guidelines, we have ensured that both male and female mice were included in all experimental groups.

### Cell culture

Primary cortical-hippocampal neuronal cultures (cortex-enriched) were prepared from pups (P1-5) as previously described [29]. 110,000 - 150,000 cells were plated on Poly-D-Lysine coated coverslips or dishes and maintained in MEM (Gibco) medium, (called “control media”) containing 0.5% Glucose (Sigma), 0.02% NaHCO3 (Roth) and 0.01% Transferrin (Merck), which was further supplemented with 5% FCS (Sigma), 0.25% L-GlutMax (100×, Gibco), 2% B27 (50×, Gibco), 1% Pen/Strep (10000 Units/ml Penicillin, 10,000 µg/ml Streptomycin, Gibco) (see **Suppl. Table 1**). After two days *in vitro* 2 mM AraC was added to the culture medium to limit glial proliferation. Primary neurons were kept at 37°C and 5% CO2. To initiate homologous recombination in neurons isolated from *Atg5*flox:*CAG-Cre*^Tmx^ animals, 0.4 μM (Z)-4-hydroxytamoxifen (Sigma) was added immediately after plating. Further, at 24 h, 0.2 µM (Z)-4-hydroxytamoxifen was added during medium supplementation providing ATG5 KO neuronal culture. An equal volume of 100% ethanol (tamoxifen vehicle) was added to WT controls.

### Cortical acute slices preparation

Mice were sacrificed at the age of 2-3 months via cervical dislocation, and brains were removed. The forebrain was cut with a vibratome (Leica) into 100 μm horizontal sections in ice-cold DPBS (Gibco, 14190144). Acute slices were then incubated in appropriate control media, control media + 10nM Rapamycin, control media without FCS (-FCS) or in EBSS (Gibco, 24010043) for either 2h or 6h at 37°C and 5% CO2 (see **Suppl. Table 1**). Following incubation cortex and hippocampus were isolated, snap-frozen and processed further for whole lysate preparation or cytosolic/synaptosome isolation (see cytosolic and synaptosome isolation from cortical acute slices) for proteomic analysis or western blotting.

### Cytosolic and synaptosome isolation from neuronal culture and cortical acute slices

Cytosolic and synaptosome fractions were isolated from postnatal cortical neuronal culture and acute brain slices (see above) following the manufacturer’s protocol (ThermoScientific™ Syn-PER™ Synaptic Protein Extraction Reagent, Catalog # 87793). Postnatal primary neurons were harvested gently via scraping on ice in Syn-PER reagent supplemented with protease/phosphatase inhibitors (10mL/g brain tissue, ThermoScientific, 15662249). Acute slices were homogenized using a Wheaton Potter-Elvehjem tissue grinder with 10 slow strokes on ice in Syn-PER reagent supplemented with protease/phosphatase inhibitors. The homogenate was then centrifuged at 1,200 x g for 10 minutes at 4°C. Then, supernatant was transferred into a new tube and centrifuged at 15,000 x g for 20 minutes at 4°C. Here, prior to centrifugation a portion of the homogenate supernatant was saved for analysis if necessary. Following centrifugation, the supernatant (cytosolic fraction) was transferred into a new tube and the pellet (synaptosomes) was re-suspended in freshly prepared Syn-PER reagent. Samples were sonicated and protein concentrations were determined using the Bradford assay (Sigma) for downstream analyses (see below). The procedure of synaptosome isolation took less than 30 min.

### SDS-PAGE and Western Blotting

For western blot analysis, primary neurons were harvested at DIV14-16 in RIPA buffer (50 [mM Tris pH 8.0, 150 mM NaCl, 1.0% IGEPAL CA-630, 0.5% Sodium deoxycholate, 0.1% SDS) containing protease inhibitor (Roche) and phosphatase inhibitor (see above) using a cell scraper. Acute brain slices were homogenized in RIPA buffer using a Wheaton Potter-Elvehjem Tissue Grinder. Isolated synaptosomes were processed further in Syn-PER reagent. Samples were sonicated and incubated on ice for 45 min prior to centrifugation at 14.000 x g for 20 min (4°C.) Supernatant concentrations were determined using Bradford assay (Sigma). Whole lysates and fractioned lysates were mixed with 4x SDS sample buffer and boiled at 95°C for 5 min. Depending on the assay, 10-20 μg protein per sample were loaded onto SDS-page gels for protein separation. Proteins were transferred onto nitrocellulose or methanol-activated PVDF membranes via full-wet transfer assay (BioRad) or semi-wet transfer. Protein transfer was confirmed by Ponceau S staining. Membranes were blocked in 5% milk or bovine serum albumin (BSA) in TBS containing 1% Tween (TBS-T) for 1h at RT followed by primary antibody (see Table S2) incubation in TBS over night at 4°C. Afterwards, membranes were washed 3 times with TBS-T for 10 min and then incubated with HRP-tagged secondary antibodies (see **Suppl. Table 3**) for 1h at RT followed by 3 washes in TBS-T for 10 min at RT. Protein levels were visualized using ECL-based autoradiography film system (Super RX-N, Fujifilm) or ChemiDocTM Imaging system (BioRad) and analyzed using Gel Analyzer plugin from ImageJ (Fiji). Protein levels were always first normalized to loading control and then to the corresponding control.

### Analysis of autophagy flux in primary cortical neurons

Cortical primary neuronal culture were incubated in osmolarity adjusted control medium with 10nM Rapamycin (selleckchem, S1039), serum free medium (-FCS) and EBSS separately and with 67nM Bafilomycin A1 (Sigma, 88899-55-2) for 6 h at 37 °C and 5% CO_2_. Cortical primary neurons were harvested gently via scraping on ice in RIPA buffer with additional of protease/phosphatase inhibitors and processed for western blot analysis (see western blotting).

### Analysis of autophagy flux in cortical acute slices

Cortical acute slices (100 μm, horizontal) were obtained as described above and incubated in control medium, control medium with rapamycin (Rapa), amino acid deficient medium (-AA), amino acid and serum deficient media (-AA/FCS), control medium without serum (-FCS) and 2mM glucose-containing medium separately and each with 400 µM chloroquine (Sigma, C6628) for 6 h at 37 °C and 5% CO_2_. Cytosolic and synaptosome fractions were isolated (as described above) and processed for western blot analysis (see western blotting).

### Immunoprecipitation

Following cytosolic and synaptosome fractionation 20 μl Dynabeads Protein G (Thermo Fischer Scientific) were coated with 2 μg antibody targeting the protein of interest and corresponding IgG (see **Suppl. Table 3**) as a negative control. Therefore, the bead’s storage solution was replaced by 100 μl PBS, and 2 μg of antibody was added. The beads were incubated with the antibody for 2–3 h at 4°C on a shaker before being washed once with 200[μl PBS to remove excess antibodies. An equal amount of protein was added to the beads coated with antibodies against the protein of interest and control IgG for overnight incubation at 4°C on a shaker. The next day, lysates were removed, and beads were washed 3 times with 100 µl Co[IP buffer (50 mM Tris-HCl; 1% IGEPAL; 100mM NaCl; 2mM MgCl_2_ in ddH_2_O (1 tablet of protease and phosphatase inhibitors/10 ml of lysis buffer) before they were dissolved in 20 μl co-IP buffer and 20 μl 4 x SDS buffer and boiled at 95°C for 5 min. Precipitation of proteins was detected via SDS-PAGE gel or proteomic analysis.

### Immunocytochemistry on cultured neurons

Directly after nutrient stress treatment neurons were fixed at DIV 12–14 in 4% paraformaldehyde/sucrose (PFA, Merck) in phosphate-buffered saline (PBS) for 15[min room temperature (RT), washed and blocked with PBS containing 5% normal goat serum (NGS, Thermo Scientific) and 0.3% Saponin (Sigma) for 1h. Neurons were incubated with primary antibodies (see Supplementary Table 3) for 1h in PBS containing 5% NGS (Thermo Scientific) and 0.3% Saponin (Roth). Coverslips were rinsed three times with PBS (10 min each) and incubated with corresponding secondary antibodies (see Supplementary Table 3) for 30 min (diluted 1:500 in PBS containing 0.3% Saponin and 5% NGS). Subsequently, coverslips were washed three times in PBS and mounted in Immu-Mount (Thermo Scientific).

Fixed neurons were imaged using either Zeiss Axiovert 200M microscope equipped with 63×/1.4 oil DIC objective and the Micro[Manager software (Micro Manager1.4, USA) or with Leica SP8 confocal microscope (Leica Microsystems) equipped with a 63×/1.32 oil DIC objective and a pulsed excitation white light laser (WLL; ∼80-ps pulse width, 80-MHz repetition rate; NKT Photonics). To assess the recruitment of autophagy-related proteins to synaptic compartments, intrasynaptic versus extrasynaptic fluorescence intensity ratios were quantified as follows. First, VGLUT1-positive puncta were identified as synaptic regions of interest (ROIs). For each ROI, mean gray values (MGVs) were measured for the protein of interest (e.g., LC3, ATG16L1, ATG5, ATG12) within the VGLUT1-defined synaptic area (intrasynaptic signal) and in adjacent non-synaptic axonal regions (extrasynaptic signal) of equivalent size. Background fluorescence was subtracted from both measurements using an average value from cell-free regions in the same field. The intrasynaptic/extrasynaptic ratio was then calculated by dividing the background-corrected MGV of the synaptic ROI by that of the extrasynaptic ROI. For quantification of Rab5b and autophagy machinery (ATG5, ATG12, ATG16L1) colocalization at synapses, colocalization was assessed by identifying VGLUT1-positive synaptic puncta and measuring the degree of spatial overlap with the indicated autophagy proteins. Pearson’s correlation coefficient was calculated after background subtraction using the Coloc2 plugin in Fiji/ImageJ.

### Electron microscopy

Cortical neurons were cultured on poly-D-lysine coated 18 mm aclar disc coverslips for 14-16 days prior to starvation treatments. After treatment they were fixed with pre-warmed fixative solution (2% glutaraldehyde, 2.5% sucrose, 3 mM CaCl2,100 mM HEPES, pH 7.4) at RT for 30 min, followed by the post-fixation at 4 °C for 30 min. Afterwards, the cells were washed with 0.1 M sodium cacodylate buffer, incubated with 1% OsO4, 1.25% sucrose, 10 mg/ml K4[Fe(CN)6]·3H2O in 0.1 M cacodylate buffer for 1 h on ice and washed three times with 0.1 M cacodylate buffer. Subsequently, the cells were dehydrated at 4 °C using ascending ethanol series (50, 70, 90, 100%, 7 min each), incubated with ascending EPON series (EPON with ethanol (1 + 1) for 1 h; EPON with ethanol (3 + 1) for 2 h; EPON alone ON; 2 × 2 h with fresh EPON at RT) and finally embedded for 48–72 h at 62 °C. Ultrathin sections of 70 nm were made using an ultramicrotome (Leica, UC7) and stained with uranyl acetate for 15 min at 37 °C and lead nitrate solution for 4 min. Electron micrographs were taken with a JEM-2100 Plus Electron Microscope (JEOL), Camera OneView 4 K 16 bit (Gatan) and software DigitalMicrograph (Gatan). The number of autophagic vesicles (AV), endosomes and multivesicular bodies (MVB) were assessed manually in a blinded fashion.

### Ciliobrevin D treatment in serum-deprived synaptosomes

Cortical acute slices (100 μm, horizontal) were obtained as described above and incubated either in control medium and control medium + 100nM Ciliobrevin D or serum free medium and serum free medium +100nM Ciliobrevin D. Following incubation for either 2hr/6hr synaptosomes were isolated as described above and samples processed for SDS-Page and western blotting.

### Transfections and live cell imaging of tagged Rab5b/ATG16L1 constructs

Cortical neurons were transfected at DIV7-9 using the optimized calcium phosphate protocol (ProFection Mammalian Transfection System-Calcium Phosphate Kit (Promega) as described previously [100]. 6 µg plasmid DNA (concentration 1 µg/µL) (Suppl. Table 4) were mixed with 12.5 µL 2M CaCl2 and 81.5 µL H_2_O and then and mixed with an equal volume of 2x HEPES buffered saline (100μl) and incubated for 20 min, allowing for precipitate formation. Meanwhile, cells were incubated in osmolarity adjusted NBA (Gibco) medium for the same time at 37°C, 5% CO2. Subsequently, precipitates were added to the cells and incubated at 37°C, 5% CO2, for 20-25 min. Finally, coverslips were washed 3x with osmolarity adjusted HBSS medium and transferred back to original maintenance medium where they were kept under 37°C, 5% CO2. Neurons 5-7 days post transfection were treated with appropriate nutrient stress conditions and then imaged at 37 °C in imaging buffer (170 mM NaCl, 3.5 mM KCl, 0.4 mM KH2PO4, 20 mM N-Tris[hydroxyl-methyl]-methyl-2-aminoethanesulphonicacid (TES), 5 mM NaHCO3, 5 mM glucose, 1.2 mM Na2SO4, 1.2 mM MgCl2, 1.3 mM CaCl2, pH 7.4) using Zeiss Axiovert 200 M microscope (Observer. Z1, Zeiss, USA) equipped with 63x/1.40 Oil DIC objective, a pE-4000 LED light source (CoolLED) and a Hammatsu Orca-Flash4.O V2 CMOS digital camera. Time-lapse images of neurons expressing mcherry-GFP-LC3 were acquired every second for 60s and puncta mCherry^+^ puncta and GFP^+^/mCherry^+^ puncta were analyzed. For neurons expressing mEmerald-ATG16 and mCherry-Rab5b images were acquired every second for 60s. Kymographs were generated using the software Kymo-Maker and analyzed by ImageJ.

### OTCs preparation and imaging

Mice were euthanized at postnatal day 6–9, cortex was collected in ice-cold HBSS (Thermo Fisher Scientific) and cut into 300-µm-thick sagittal sections. Sections were washed three times in pre-warmed (37 °C) HBSS. Slices were then transferred onto membrane inserts (Merck) with pre-warmed OTC MEM-based (Sigma) medium containing 0.00125% ascorbic acid (Roth), 10 mM D-glucose (Sigma), 1 mM GlutaMAX (100× Gibco), 20% (vol/vol) horse serum (Gibco), 0.01 mg ml^−1^ insulin (Thermo Fisher Scientific), 14.4 mM NaCl (Roth), 1% penicillin–streptomycin (Thermo Fisher Scientific). Medium was replaced every second day. Viral transduction was done on DIV1 by adding 1µl of hSyn1-*Syph*-BFP2 and 1µl of hSyn1-mcherry-EGFP-*mMap1lc3b* (Suppl. Table 5) on top of each slice. Live cell imaging was performed at DIV 21. 2 hr prior to imaging media was replaced with either fresh control medium or medium lacking serum. OTCs were places on µ-dish polymer coverslip (Ibidi), containing the respective media, imaged on Lecia Stellaris 5 equipped with an Apo x63/1.32 FLYC CORR CS2 objective and a continuous excitation white light laser. OTCs were imaged at 2,048×2,048 pixels with a bidirectional recording. Images were analysed using Fiji (ImageJ) mCherry^+^/eGFP^+^ puncta between the sizes 0.5-2.5µm^2^ were counted within the BFP^+^ synapses.

### Calcium imaging in primary neurons

Primary cortical neurons were transduced with hSyn[jGCaMP7f (see Supplementary Table 5) at DIV7 and maintained at 37°C, 5% CO2 until DIV15-16 when appropriate nutrient stress medium was applied for 2 hr prior to imaging. Live-cell imaging and stimulation was performed using Zeiss Axiovert 200M microscope (Observer. Z1, Zeiss, USA) equipped with 10×/0.3 EC Plan[Neofluar objective; a pE-4000 LED light source (CoolLED), and a Hamamatsu Orca-Flash4.O V2 CMOS digital camera. Neurons were imaged in an osmolarity adjusted imaging buffer described previously (Kononenko et al, 2013; 120 mM NaCl, 3.5 mM KCl, 0.4 mM KH_2_PO_4_, 20 mM TES, 5 mM NaHCO_3_, 5 mM glucose, 1.2 mM Na_2_SO_4_, 1.2 mM MgCl_2_, 1.3 mM CaCl_2_, pH 7.4), and time-lapse images were taken within a 1 s interval. For electrical field stimulation, neurons were stimulated 4x with 100 action potentials (100 AP) in a 3 s interval at 100 Hz using an RC-47FSLP stimulation chamber (Warner Instruments). For analysis, neuronal cell bodies were chosen as ROI, and the GCaMP7f fluorescence response to the stimulus was plotted over time using Image J (Plot Z axis Profile tool). Baseline fluorescence value was subtracted after background subtraction, and the traces were normalized to the first peak to visualize the facilitation.

### FRAP imaging

Live-cell imaging of cortical neurons expressing eGFP–ATG5 was performed using Leica Stellaris 5 equipped with an Apo x63/1.32 FLYC CORR CS2 objective and a continuous excitation white light laser, equipped with a live-cell incubation chamber. Puncta along neurites were photobleached using a focused 488 nm laser pulse (100% intensity, 1s duration), followed by time-lapse imaging at 0, 1, 30, and 60 s post-bleaching. Fluorescence intensity was quantified within the bleached ROI and normalized to pre-bleach values. Background signal was subtracted using a nearby ROI outside the cell. Fluorescence recovery was plotted over time, and area under the curve (AUC) was calculated to compare protein mobility across conditions. FRAP measurements were performed on neuritic puncta and were not explicitly restricted to VGLUT1-marked synapses; thus, stabilization at synapses is inferred from complementary synaptic recruitment and fractionation data.

### Dietary interventions in 8-week-old mice

Dietary preconditioning of male C57BL6N was approved by the state authorities (Landesamt für Natur, Umwelt und Verbraucherschutz Nordrhein-Westfalen (LANUV) AZ2019.A345) and performed as previously described in [58]. Briefly, male C57BL6N mice aged 8 weeks were allocated randomly to their specific dietary regimen and transferred to a fresh cage at beginning of dietary intervention. FMD, S-AA, BCAA and CR (see Supplementary Table 6 for diet composition) were performed for 14 days. During dietary preconditioning mice had *ad libitum* access to water and they consumed all supplied diets revealing no signs of food aversion. To assess weight loss mice were weighed weekly and no increase in morbidity or mortality was observed during any of the dietary interventions. Mice were euthanized using cardiac blood drawing followed by PBS-perfusion and brains were isolated and immediately snap frozen in liquid nitrogen for further analysis.

### Proteomics of synaptosomes from cultured WT vs ATG5KO neurons

Total proteome analysis from WT and ATG5KO synaptosomes were isolated as described above. Samples were homogenized and lysed in 8 M urea followed by reduction (5 mM dithiothreitol (DTT)) and alkylation of cysteines (40 mM chloroacetamide). Samples were digested using LysC for 4 h followed by dilution of urea and overnight digestion using trypsin (both with a 1:75 ratio). Afterwards, samples were acidified and clean-up was performed using mixed-mode StageTips. Samples were analyzed by the CECAD Proteomics Facility on an Orbitrap Exploris 480 mass spectrometer (Thermo Scientific, granted by the Deutsche Forschungsgemeinschaft INST 216/1163-1 FUGG) coupled to an Evosep ONE. The Evosep was run with its Whisper Zoom 20 SPD gradient using a PepSep column with the recommended parameters (15 cm length, 75 µm inner diameter, 1.9 µm diameter C18 material, Bruker) and a 10 µm glass emitter (Evosep). The mass spectrometer was operated using a WHISH-DIA approach [101]. MS1 spectra were acquired between 400 and 1000 m/z with a resolution of 120k, MS2 spectra were acquired in the same range at 60k resolution in 25 m/z windows, resulting in 24 scans total. Fragments were acquired in a range of 250 to 1500 m/z with a normalized AGC target of 1000% and 30% normalized HCD collision energy. Every 6 scans, an MS1 scan was inserted. Samples were analyzed in Spectronaut 19 (Biognosys) after conversion into htrms format using a murine SwissProt database (UP0589, downloaded 15/01/2024). Standard settings were used but quantification was calculated on MS1 level. Follow-up analyses were performed in Perseus 1.6.15 with data cleanup and filtering for data completeness in at least one group. Afterwards, FDR-controlled T-tests were performed, followed by enrichment analyses using GO and KEGG annotations provided by the SynGo website [31].

### Proteomics of synaptosomes isolated from acute brain slices

Synaptosomes were isolated as described above. For proteomic analysis of synaptosomes, proteins were isolated with 8M urea in 100mM Tris, pH 7.5. Protein concentration was determined with BCA and 10µg were subjected sequentially to reduction with 10mM DTT for 15min, alkylation with 55mM CAA for 30min and digestion with LysC (1:100 enzyme:substrate ratio) for 2hr. Samples were diluted with 100mM Tris pH 7.5 to a final concentration of 2M urea and further digested with Trypsin (1:100 enzyme:substrate ratio) over night at room temperature. Peptides were desalted and stored on SDB-RPS stage tips. For measurement, peptides were eluted from stage tips, dried in a speed vac and resuspended in 100µl resuspension buffer (5% formic acid (FA) and 2% acetonitrile (ACN)). Samples were measured on a Bruker TimsTOF pro2 coupled to a Bruker NanoElute2 HPLC using a 25cm Bruker PepSep chromatographic column in a DIA-PASEF mode. Each sample was measured with a 60min reverse phase gradient and 300ng were injected for measurement, respectively. Resulting raw data were converted into “.dia” format in DIA-NN (v.1.8.1) [102] followed by a library-free analysis using the UniProt mouse protein database (release 2023-05) including canonical, isoforms and Trembl hits. DIA-NN settings were left by default with the following exceptions: (1) protein inference set to “Isoform IDs”, (2) Neural network classifier set to “Double-pass mode”, (3) Heuristic protein inference unchecked, (4) Additional options include command “report-lib-info”. The DIA-NN output was further processed with an in-house R script, to apply Max-LFQ values for quantification of protein intensities. Resulting table was analysed with Perseus (v.2.0.11) [103], InstantClue (v.0.12.0) [104] and RStudio.

### Proteomics from ATG16L1 pulldown from cytosolic and synaptosome fractions isolated from acute brain slices

ATG16L1 precipitations were washed thoroughly and digested with trypsin over night. Resulting peptides were desalted and stored on SDB-RPS stage tips. For measurement, peptides were eluted, dried in a vacuum concentrator and resuspended in 20µl resuspension buffer. Samples were measured on a Thermo Q Exactive Plus Orbitrap instrument coupled to a Thermo Ultimate3000 HPLC with a 30min reverse phase gradient in a DIA mode with staggered isolation windows of 24m/z. Resulting raw files were demultiplexed and converted into “.mzml” format using the MSconvertGUI tool (ProteoWizzard) and analysed in DIA-NN (v.1.8.1). DIA-NN settings were left by default with the following exceptions: (1) protein inference set to “Isoform IDs”, (2) Neural network classifier set to “Double-pass mode”, (3) Heuristic protein inference unchecked. The DIA-NN output was further processed with an in-house R script, to apply Max-LFQ values for quantification of protein intensities. Resulting table was analysed with Perseus (v.2.0.11) [103], InstantClue (v.0.12.0) [104] and RStudio.

### Proteomics of mouse brains after 2 weeks of dietary innervations

Proteins were extracted in 4% SDS/PBS lysis buffer. Protein concentration was determined with a bicinchoninic acid (BCA) assay (Pierce). 250 µg of each lysate were reduced with 5mM Tris(2-carboxyethyl) phosphine (TCEP) and alkylated with 15mM chloroacetamide (CAA) for 15min at 75°C. Samples were digested following the standard SP3 protocol [105]. Briefly, 62.5 μl of washed magnetic SP3 beads (Sera-Mag, Sigma Aldrich) were added to each sample, an equal volume of 100% acetonitrile (ACN) was added to achieve a final concentration of 50% ACN. After incubation for 8 min, the supernatant was discarded, and SP3 beads were washed twice with 70% EtOH and once with 100% ACN. Digestion was performed by resuspending beads in 250 µL of 50 mM Triethylammoniumbicarbonat (TEAB) buffer supplemented with the endoproteinases LysC (1:200 w/w, KPL, Denmark) and Trypsin (1:100 w/w, Serva, Germany). Samples were incubated overnight at 37°C on a thermomixer (550 rpm), acidified by addition of 100 μL 0.1% FA, followed by clean-up with SepPak C18 cartridges (Waters, USA). Peptides were reconstituted in 2% ACN, 5% FA and 200 ng peptides were loaded on EvoTips. Using a 30 sample per day (SPD) gradient on an EvoSep One system coupled to a 15 cm Performance column (EV1137 ReproSil-Pur C18, 1.5 µm beads, 15 cm x 150 µm, Evosep, Denmark) samples were measured on a Thermo Eclipse Orbitrap instrument (Thermo-Fisher Scientific, Waltham, USA) with a staggered DIA method with 30 x 20 mz windows ranging from 400 to 1000 m/z. Resulting raw files were demultiplexed and converted into “.mzml” format using the MSconvertGUI tool (ProteoWizzard) and analysed with a library-free search in Spectronaut (version 19.0.240606, Biognosys, Switzerland) using the canonical UniProt mouse protein database (downloaded 15.01.2025).

### Statistical analysis

Sample sizes were not chosen based on pre[specified effect size. Instead, multiple independent experiments were carried out using several samples replicates as detailed in the figure legends. For all experiments, there was enough statistical power to detect the corresponding effect size. Statistical analyses were done on cell values (indicated by data points) from at least three independent experiments (indicated by “n”, biological replicates). MS Excel (Microsoft, USA) and GraphPad Prism version 9 (GraphPad Software, Inc., USA) were used for statistical analysis and result illustration (unless otherwise stated). Statistical significance between two groups for normally distributed non[normalized data was evaluated with a two-tailed unpaired Student’s t-test. Statistical difference between more than two groups and two conditions was evaluated using Two-Way ANOVA (Holm–Sidak post hoc test or Benjamini, Krieger and Yekutieli post hoc test were used to determine the statistical significance between the groups). Statistical difference between more than two groups was calculated using One-way ANOVA with Holm–Sidak post hoc test. A normality test using GraphPad Prism version 9 (Shapiro-Wilk test) was performed to determine the parametric and nonparametric statistical tests used for analysis (for data sets with > 8 data points). Data were excluded only when sample quality was not optimal and/or due to technical artifacts (e.g., imaging failure, tissue damage and/or due to re-genotyping results. Predefined quality criteria were: in living samples-vacuolization or other signs of cellular degeneration, normal cell morphology and protein transport (which is usually hampered in unhealthy cells) and image streams that are in focus; proper sample preparation and mounting in fixed samples.

Because the number of quantified proteins in synaptosomal preparations was limited and the effect sizes modest, global Benjamini–Hochberg false discovery rate (FDR) correction was found to be overly conservative, eliminating biologically meaningful signals. Importantly, the assumptions underlying FDR procedures (large test numbers and independence of features) are not fully met in our dataset, where synaptic proteins are highly co-regulated. Therefore, and consistent with common practice in discovery-driven MS studies with limited depth, statistical interpretation was based on moderated p-values together with effect-size thresholds (e.g., log2 fold change > 0.5). These criteria have been widely used in proteomics to balance sensitivity and specificity in small-sample datasets.

Significant differences were accepted at p<0.05 indicated by asterisks: *p<0.05; **p<0.005; ***p<0.0005.

## Resource availability

### Lead contact

Further information and requests for resources and reagents should be directed to and will be fulfilled by the lead contact, Natalia Kononenko.

### Data and code availability

- Proteome data of all experiments are deposited in the database PRIDE and publicly accessible after publishing.
- Source data are provided with this paper.
- Datasets supporting the current study will be shared by the lead contact upon request.

## Acknowledgements

We thank R. Renn and Dr. M. Schröter for their expert assistance. We are indebted to Dr. C. Jüngst (CECAD Imaging Facility), Dr. S. Müller and Dr. J.-W. Lackmann (CECAD Proteomic Facility) for their help and expert assistance. The work of NLK is funded by the Deutsche Forschungsgemeinschaft (DFG, German Research Foundation): EXC 2030–390661388, KO 5091/4-1, DFG-431549029–SFB 1451 and DFG-411422114-GRK 2550.

## Author contributions

Conceptualization: MO, NLK Methodology: MO, NLK Investigation: MO, LI, AH, SM, FT Visualization: MO, LI, AH, SM, FT Supervision: RUM, MK, NLK Writing—original draft: NLK Writing—review & editing: NLK

