## Supplementary Material for "Nutrient stress activates Rab5b-mediated autophagy to remodel the synaptic proteome"

\# shared first author

**Supplementary Table 1 – Cell culture media compositions and starvation media**

| **Medium** | **Base Medium** | **Glucose** | **NaHCO₃** | **NaCl** | **CaCl₂** | **MgSO₄·7H₂O** | **KCl** | **NaH₂PO₄·H₂O** | **Transferrin** | **L-GlutaMax (100×)** | **B27 (50×)** | **Pen/Strep** | **FBS** |
| --- | --- | --- | --- | --- | --- | --- | --- | --- | --- | --- | --- | --- | --- |
| Control media | MEM (Gibco) | 10mM | 26.19 mM | 117.24 mM | 1.8 mM | 0.81 mM | 5.3 mM | 0.02% | 0.01% (Merck) | 0.25% (Gibco) | 2% (Gibco) | 1% (Gibco) | 5% (Sigma) |
| EBSS (Gibco) | EBSS (Gibco) | 5.5 mM | 26.19 mM | 117.24 mM | 1.8 mM | 0.81 mM | 5.5 mM | 1.01 mM | – | – | – | – | – |
| Serum-Free DMEM-MEM | MEM (Gibco) | 10mM | 26.19 mM | 117.24 mM | 1.8 mM | 0.81 mM | 5.3 mM | 0.02% | 0.01% (Merck) | 0.25% (Gibco) | 2% (Gibco) | 1% (Gibco) | – |
| AA-Free DMEM (w/ FBS) | AA-free MEM (Genaxoon) | 10mM | 44.04mM | 110.3mM | 1.8 mM | 0.84mM | 5.3 mM | 0.9 mM | 0.01% (Merck) | 0.25% (Gibco) | 2% (Gibco) | 1% (Gibco) | 5% (Sigma) |
| AA & Serum-free DMEM | AA-free MEM (Genaxoon) | 10mM | 44.04mM | 110.3 mM | 1.8 mM | 0.84mM | 5.3 mM | 0.9 mM | 0.01% (Merck) | 0.25% (Gibco) | 2% (Gibco) | 1% (Gibco) | - |
| 2mM Glucose | DMEM low glucose (Gibco) | 2mM | - | 110.3 mM | 1.8 mM | 0.81 mM | 5.3 mM | 0.9 mM | 0.01% (Merck) | 0.25% (Gibco) | 2% (Gibco) | 1% (Gibco) | 5%(Sigma) |
| Control OTC Media | MEM (Sigma) | 10 mM | - | 86 mM | 1.4 mM | 0.616 mM | 4.1 mM | 0.9 mM | - | 1 mM | - | 1% | 20% (heat inactivated horse serum) |
| Serum Free OTC | MEM (Sigma) | 10 mM | - | 86 mM | 1.4 mM | 0.616 mM | 4.1 mM | 0.9 mM | - | 1 mM | - | 1% | - |

**Supplementary Table 2 – Primers used for genotyping**

| **Gene** | **Sequence (5’- 3’)** |
| --- | --- |
| *Atg5* forward 1 | GAA TAT GAA GCC ACA CCC CTG AAA TG |
| *Atg5* forward 2 | ACA ACG TCG AGC ACG CTG GCG AAG G |
| *Atg5* reverse | GTA CTG CAT AAT GGT TTA ACT CTT GC |
| Cre 1 | GAA CCT GAT GGA CAT GTT CAG G |
| Cre 2 | AGT GCG TTC GAA CGC TAG AGC CTG T |
| Cre 3 | TTA CGT CCA TCG TGG ACA |
| Cre 4 | TGG GCT GGG TGT TAG C |

**Supplementary Table 3 – List of primary and secondary antibodies**

| Antibody | Used concentration | | | Manufacturer | Product ID |
| --- | --- | --- | --- | --- | --- |
| Antibody | Dilution WB | Dilution IF | Concentration in IP |  |  |
| P62 | 1:1000 | - | - | Progen | GP62-C |
| LC3B | 1:1000 | 1:500 | - | Novus  Biologicals | NB600-  1384SS |
| ACTB | 1:2000 | - | - | Sigma | A5441 |
| VGLUT1 | - | 1:1000 | - | SySy | 135304 |
| ATG5 | - | 1:500 | - | Novus Biologicals | NBP1-76992 |
| ATG5 | 1:1000 |  | 2µg | Abcam | ab108327 |
| ATG12 | 1:000 | 1:500 | 2µg | Proteintech | 11122-1-AP |
| ATG16l1 | - | 1:500 | - | Proteintech | 67943-1-Ig |
| ATG16l1 | 1:1000 | - | - | MBL |  |
| PSD95 | 1:1000 | - | - | Sysy | 124011 |
| RAB5b | - | 1:500 | - | Proteintech | 27403-1-AP |
| ATG16L1 | 1:1000 | - | 2µg | Cell Signaling | 8089 |
| Phospho-S6 (235/236) | 1:1000 | - | - | Cell Signaling | 2211S |
| Rabbit IgG | - | - | 2µg | Cell Signaling | 2729 |
| Alexa Fluor™ 488 Donkey Anti-Rabbit IgG |  | 1:500 |  | Invitrogen | A32790 |
| Alexa Fluor 568 Goat Anti-Guinea Pig IgG |  | 1:500 |  | Life Technologies GmbH | A11075 |
| Alexa Fluor™ 488 Donkey Anti-Mouse IgG |  | 1:500 |  | Life Technologies GmbH | A32766 |
| Alexa Fluor 488 Goat Anti-Rabbit IgG |  | 1:500 |  | Life Technologies GmbH | A11034 |
| Alexa Fluor 568 Goat Anti-Chicken IgG |  | 1:500 |  | Life Technologies GmbH | A11041 |
| Alexa Fluor 647 Goat Anti-Mouse IgG |  | 1:500 |  | Life Technologies GmbH | A21236 |
| Goat-Anti-Rabbit-HRP | 1:10,000 |  |  | Dianova GmbH | 111-035-003 |
| Goat-Anti-Mouse-HRP | 1:10,000 |  |  | Dianova GmbH | 115-035-003 |
| TrueBlot Rabbit IgG-HRP | 1:1000 |  |  | Rockland | **ABIN1589974** |

**Supplementary Table 4 – Plasmid list**

| Plasmid (source gene) | Manufacturer | Identifier |
| --- | --- | --- |
| mCherry-GFP-LC3 | Addgene | #170446 |
| mEmerald-ATG16L1 | Addgene | #54007 |
| mCherry-Rab5b | Addgene | #49201 |
| pEGFP-C1-hAPG5 (ATG5-GFP) | Addgene | #22952 |

**Supplementary table 5 – AAV list**

| Name in the paper | Full description | Manufacturer | Identifier |
| --- | --- | --- | --- |
| mSyp-BFP2 | AAV9S-Syn1>m*Syp*-TagBFP2-WPRE | Vector Builder | 231025-1155ejb |
| GCaMP7f | AAV9-pGP-Syn1>jGCaMP7f-WPRE (AAV1) | Addgene | 104488 |
| mCherry-eGFP-LC3B | AAV9S-SYN1>mCherry/EGFP/m*Map1lc3b*-WPRE | Vector Builder | VB240702-1321yqg |

**Supplementary table 6 – Chow information for dietary interventions**

| Diet details | Diet details |
| --- | --- |
| FMD | Day 1: S5159-E736, ssniff GmbH, Soest, Germany  Day 2/3: S5159-E738, ssniff GmbH, Soest, Germany  Day 4-7: V1554-330 ssniff GmbH, Soest, Germany |
| BCAA | S5159-E724, ssniff GmbH, Soest, Germany |
| S-AA | S5159-E720, ssniff GmbH, Soest, Germany |
| CR | 60% of daily intake (3g/day), V1554-330 ssniff GmbH, Soest, Germany |
| *ad-libitum* | V1554-330 ssniff GmbH, Soest, Germany |

**Supplementary Figures**

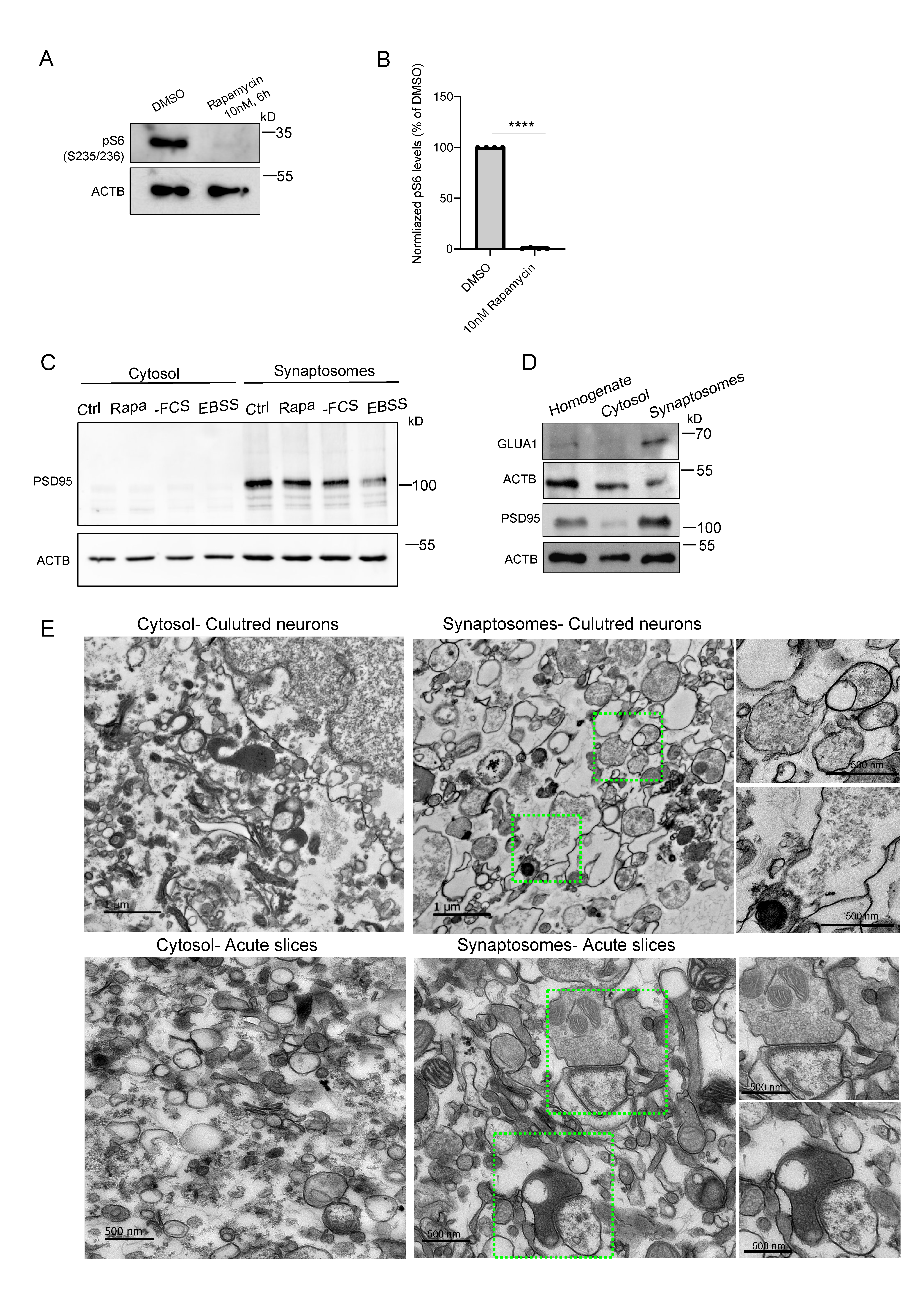

**Supplementary Figure S1. Validation of mTORC1 inhibition and quality control of cytosolic and synaptosomal fractions.**

(**A,B**) Immunoblot analysis of phospho-S6 (Ser235/236) levels in total cortical lysates from neurons treated with DMSO (control) or 10 nM rapamycin for 6 h. Values are normalized to ACTB and presented relative to DMSO controls. n=4 biological replicates.

(**C**) Immunoblot analysis of cytosolic and synaptosome fractions isolated from acute cortical slices incubated in either control maintenance medium (Ctrl), serum-free medium (-FCS), EBSS and/or control medium supplemented with 10nM rapamycin. Blots were probed for ACTB (loading control) and PSD95 (postsynaptic marker) to assess synaptic enrichment across conditions. Representative example from n=3 biological replicates.

(**D**). Immunoblot validation of subcellular fractionation from acute cortical slices: homogenate, cytosolic fraction, and synaptosomal fraction. GLUA1 and PSD95 are strongly enriched in synaptosomes, confirming high-quality fractionation.

(**E**) Transmission electron microscopy (TEM) images demonstrating the ultrastructure of cytosolic and synaptosomal preparations derived from cultured neurons and acute cortical slices. Synaptosome preparations contain intact pre- and postsynaptic profiles with preserved vesicle clusters, whereas cytosolic fractions lack membrane-enclosed structures. Scale bars: 1 µm (for primary cultures) and 500 nm (for acute brain slices).

Data represent mean ± SEM. ****p < 0.0001 by unpaired Student’s T-test.

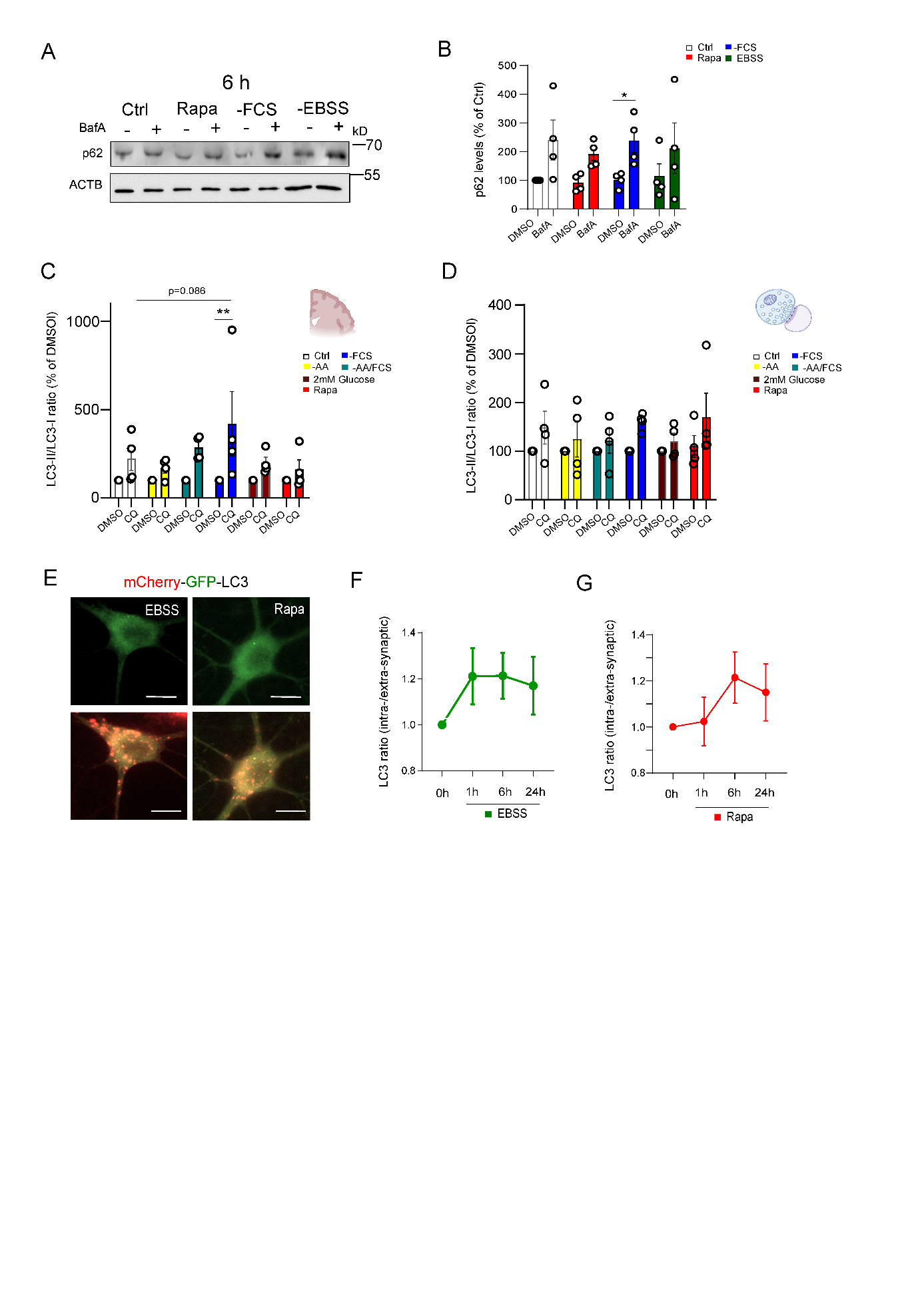

**Supplementary Figure S2. Distinct autophagy responses to serum deprivation, EBSS, and rapamycin in cortical neurons.**

(**A**) Representative immunoblot of p62/SQSTM1 in primary cortical neurons treated for 6 h with DMSO (Ctrl), 10 nM rapamycin, serum-free medium (–FCS) or EBSS, with or without 67nM BafA. ACTB serves as loading control.

(**B**) Quantification of p62 levels normalized to ACTB. A significant accumulation of p62 was observed upon –FCS treatment, indicating increased autophagic flux. n = 4 biological replicates.

(**C**) Quantification of LC3-II/LC3-I ratios in the cytoplasmic fraction of acute cortical slices. LC3 ratios were normalized to DMSO-treated controls. CQ treatment significantly increased LC3-II/LC3-I ratios under –FCS, suggesting enhanced autophagic flux. n = 3 biological replicates.

(**D**) Quantification of LC3-II/LC3-I ratios in synaptosomal fractions of acute cortical slices. LC3 ratios were normalized to DMSO-treated controls. Data from n = 4 biological experiments.

(**E**) Representative live-cell images of mCherry–GFP–LC3-expressing neurons following 6h treatment with either EBSS or rapamycin for the indicated durations.

(**F,G**) Intra-synaptic/extra-synaptic LC3 signal ratios under increasing durations of rapamycin (F) or EBSS (G) treatment (0 h, 1 h, 6 h, 24 h). Only modest increases were observed, indicating limited synaptic autophagy induction under these conditions. Data from n = 4 independent experiments.

Data represent mean ± SEM. *p < 0.05, **p < 0.005; statistical analysis by two-way ANOVA with Sidak post hoc test.

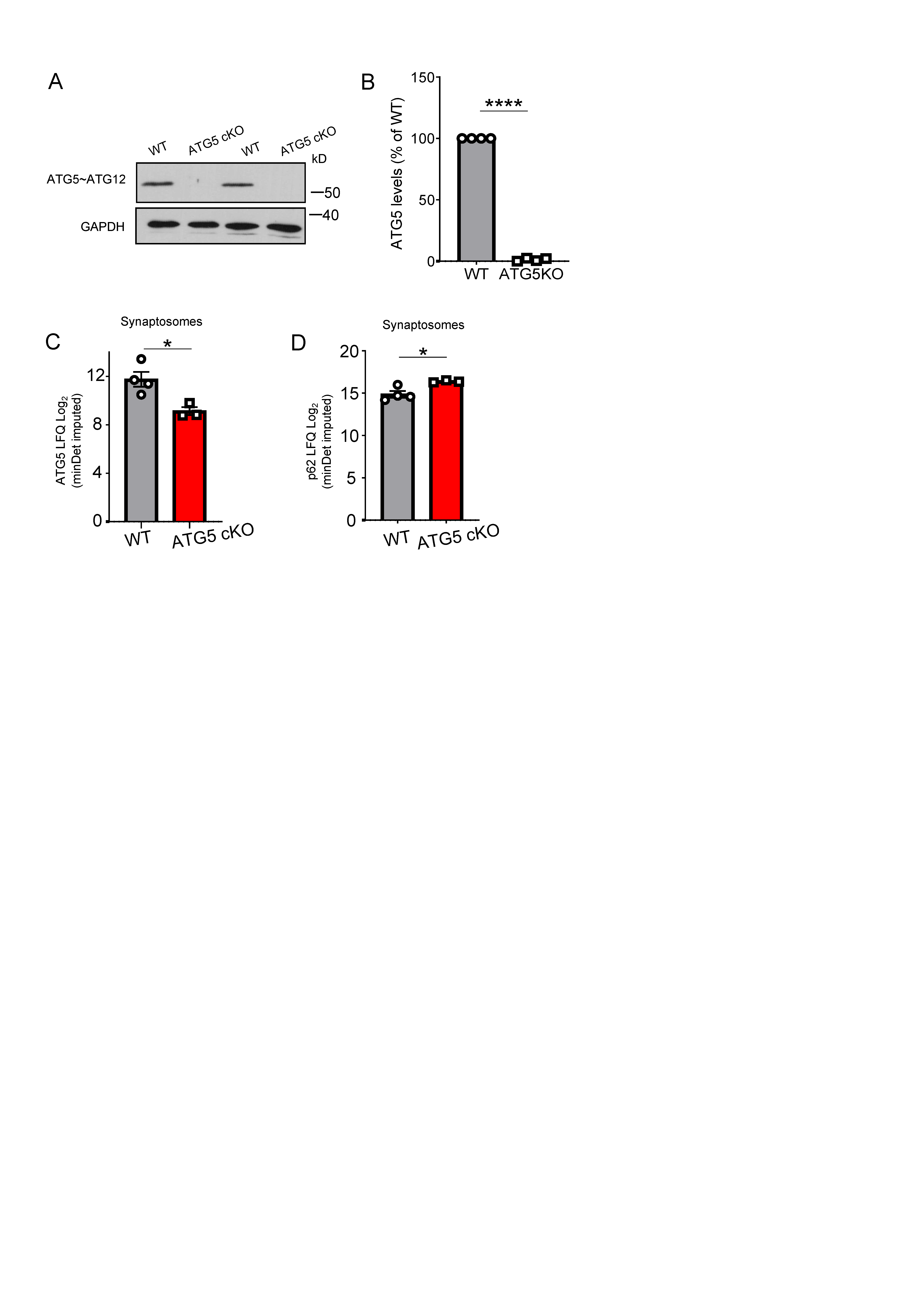

**Supplementary Figure S3. Validation of ATG5 deletion in primary neurons and synaptosomes isolated from ATG5 cKO mice.**

(**A,B**) Immunoblot analysis confirms efficient ATG5 deletion in primary ATG5 cKO neurons. Representative blots probed for the ATG5~ATG12 conjugate and GAPDH (loading control). n=4 biological replicates.

(**C**) Quantitative proteomic analysis (LFQ values) of ATG5 levels in synaptosomes isolated from WT and ATG5 cKO cortical neurons shows a significant reduction of ATG5 in KO preparations.
(**D**) Quantitative proteomic analysis (LFQ values) of p62/SQSTM1 levels in WT and ATG5 cKO synaptosomes under basal (unstimulated) conditions reveals significant accumulation of p62 in the absence of ATG5.

Data represent mean ± SEM. *p < 0.05, **** p<0.00005; statistical analysis by unpaired Student’s T-test.

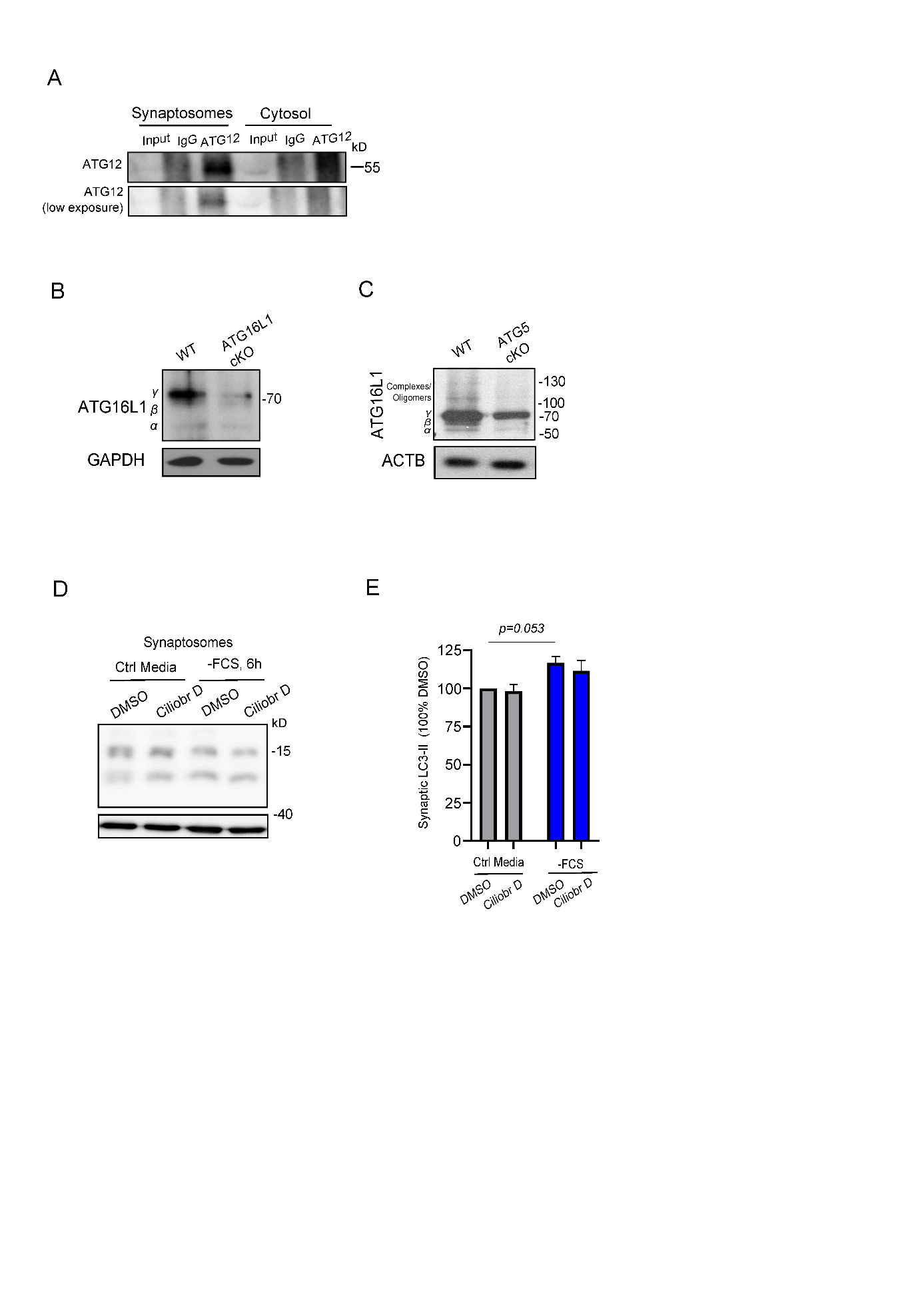

**Supplementary Figure S4. ATG16L1 isoform expression and dynein-dependent regulation of synaptic autophagy.**

(**A**) Immunoprecipitation of endogenous ATG12 from cytosolic and synaptosomal fractions of acute cortical slices followed by immunoblotting.

(**B, C**) Immunoblot analysis of ATG16L1 isoforms (α, β, γ) in the cortex of WT and ATG16L1 cKO (B) and/or hippocampus of ATG5 cKO (C) confirms the specificity of the ATG16L1 antibody, as signal is reduced in KO samples. ACTB serves as loading control.
(**D**) Immunoblot analysis of cortical synaptosomes treated with DMSO or ciliobrevin D (100 µM, 6h), with or without prior serum withdrawal (–FCS).
(**E**) Quantification of synaptosomal LC3-II levels in the same conditions as in (D).

Data represent mean ± SEM. statistical analysis by two-way ANOVA with Benjamini, Krieger, and Yekutieli post hoc test.

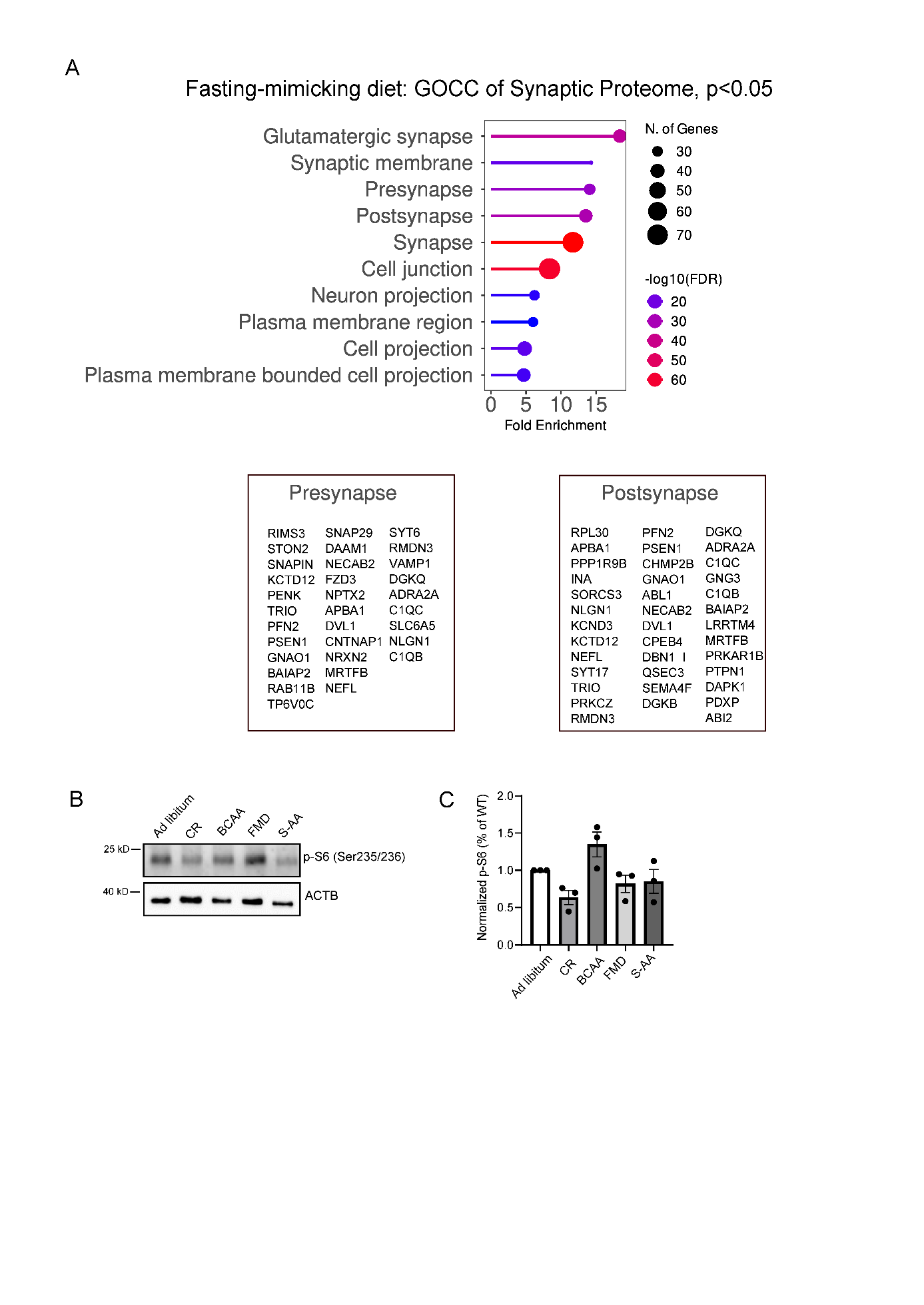

**Supplementary Figure S5. Fasting-mimicking diet remodels the synaptic proteome without significantly altering cortical phopsho-S6 activity.**

(**A**) Gene Ontology Cellular Component (GOCC) analysis of synaptic proteins significantly regulated in cortex of mice subjected to 14 days of fasting-mimicking diet (FMD). Enrichment in synaptic categories indicates selective remodeling of synaptic compartments in response to nutrient restriction.

(**B**) Representative immunoblot of phospho-S6 (Ser235/236) and ACTB in cortical lysates from mice under *ad libitum* feeding, caloric restriction (CR), branched-chain amino acid restriction (BCAA), fasting-mimicking diet (FMD), or sulfur-containing amino acid depletion (S-AA).
(**C**) Quantification of p-S6 (Ser235/236) levels normalized to ACTB. Protein levels of *ad libitum* controls were normalized to 1. n=3 mice per group; data represent mean ± SEM.
